# Lactational transfer of synthetic oxytocin and sex-specific suppression of neurodevelopmental risk gene networks in the rat prefrontal cortex

**DOI:** 10.64898/2026.05.19.726042

**Authors:** Gokhan Serce, Tusar Giri, Minsoo Son, Rencheng Wang, Marie Laury, Rebecca George, Katherine B. McCullough, Susan E. Maloney, Eric Tycksen, Young Ah Goo, Arvind Palanisamy

## Abstract

Autism spectrum disorder and related neurodevelopmental disorders (NDDs) affect males at two-to-four times the rate of females, yet the environmental contributors to this sex-differentiated vulnerability remain poorly understood. Here, using a clinically translatable postpartum rat model, we show that synthetic oxytocin (OT) administered for postpartum hemorrhage (PPH) prophylaxis (‘PPH-OT’), a universally administered obstetric medication, undergoes lactational transfer, confirmed by stable isotope-labeled OT and LC-MS/MS, without altering maternal behavior or neonatal cortical OT receptor expression. In offspring, PPH-OT altered ultrasonic vocalizations during dyadic interaction and caused a male-specific reduction in social approach, without affecting other behavioral domains. Sex-stratified RNA-sequencing of the medial prefrontal cortex (mPFC) revealed pronounced transcriptional dimorphism: males showed 1,732 differentially expressed genes versus 693 in females, and a sex-by-treatment interaction contrast identified 1,229 formally sex-differentiated genes. Male mPFC downregulated genes were significantly enriched for high-confidence NDD risk genes across three independent databases — SFARI score-1 (highest confidence autism genes; OR = 4.56, p = 0.015), DBD autism (OR = 10.34, p = 7.3×10⁻¹²), and SysNDD autosomal dominant (OR = 9.78, p = 8.4×10⁻⁷) — including core regulators of cortical circuit assembly (*Grin2a*, *Grin2b*, *Reln*, *Tbr1*, *Mef2c*, and *Tcf4*), an enrichment pattern largely restricted to males. Hypothalamic transcriptional changes were pronounced but showed no enrichment for NDD risk genes, establishing mPFC specificity. These findings identify PPH-OT as a candidate environmental modifier of male neurodevelopmental vulnerability and provide a pressing rationale for prospective investigation of neurodevelopmental outcomes in PPH-OT-exposed children.

## INTRODUCTION

Autism spectrum disorder and related neurodevelopmental disorders (NDDs) affect males at two-to-four times the rate of females, one of the most replicated and least explained observations in psychiatric epidemiology.^1–3^ Despite decades of research, the environmental contributors to this sex-differential vulnerability remain poorly understood. Oxytocinergic signaling is fundamental to maternal-neonatal bonding,^4–6^ early social brain development,^7–9^ and the ontogeny of prefrontal circuits that govern sociality throughout life.^10–13^ Therefore, perinatal disruption of such signaling is a plausible but largely unexamined candidate. Synthetic oxytocin administered for postpartum hemorrhage prophylaxis (PPH-OT) is one of the most universally administered obstetric drugs worldwide,^14–16^ delivered after both vaginal and cesarean deliveries at doses substantially exceeding physiological surges of endogenous OT during unmedicated childbirth.^17^ Whether this near-universal pharmacological exposure reaches the developing offspring through breastmilk and influences early brain development has never been systematically examined.

Emerging but conflicting observational studies suggest that perinatal OT exposure may have neurodevelopmental consequences for the offspring, including controversial associations with autism spectrum and other NDDs.^18–21^ These studies are difficult to interpret for two reasons: the inability to account for the numerous confounding variables at delivery that independently affect child neurodevelopment, and the ethical impossibility of a randomized trial given the near-universal mandated use of PPH-OT. Critically, none of these observational studies specifically examine the potential contribution from PPH-OT, a postpartum intervention administered after birth. The biological question of whether maternally administered OT can reach the neonate via lactation has received no attention so far. Despite evidence that exogenous OT can be secreted in breast milk,^22^ and absorbed via the neonatal gastrointestinal tract,^23^ the impact of PPH-OT on the developing brain remains largely unknown.

Animal studies have attempted to address the impact of perinatal OT manipulation on maternal and neonatal behaviors. These studies, including work from our laboratory, focus either on the impact of disrupted OT signaling on maternal behaviors,^6, 24–26^ the adverse neurobehavioral consequences of direct OT manipulation in the neonate,^27^ or the neurodevelopmental effects of OT exposure prior to birth.^28–30^ None of these studies mimic the scenario of postnatal exposure from maternally administered exogenous OT such as during PPH-OT. A major technical challenge has been replicating the precise intravenous infusion of PPH-OT in a postpartum dam without compromising maternal nurturing behaviors (e.g., interference from a tethered infusion system). Our laboratory has developed and validated a microprocessor-controlled implantable pump system for peripartum OT delivery,^30–32^ which we adapted here to deliver high-dose intravenous OT after delivery of the pups, simulating PPH-OT without interrupting mother-infant interactions. Here, we demonstrate that PPH-OT undergoes lactational transfer to the neonate, and characterize its sex-specific impact on neurobehavior of the offspring and its region-specific consequences for the transcriptome of the developing brain.

## METHODS

Timed-pregnant CD^®^ IGS rats (Crl:CD[SD] outbred; Charles River Laboratories) were used for the study. All animal experiments and methods reported here were approved by the Institutional Animal Care and Use Committee at Washington University in St. Louis (#23-0021) and conducted in compliance with institutional and ARRIVE 2.0 guidelines. Experiments were done in multiple cohorts over an approximately 2-year period (**Supplementary Table S1**).

### 1. iPrecio pump implantation and PPH-OT infusion

Implantation of saline-primed iPrecio pump and jugular vein cannulation was performed on gestational days (GD)16 as described by us previously with some modifications.^30–32^ These modifications included: (1) replacement of saline immediately after birth of pups with oxytocin (600 mcg/mL) in the PPH-OT group, and (ii) mock surgery without pump implantation or jugular vein cannulation in the control group. The iPrecio pump was preprogrammed to deliver oxytocin (600 μg/mL) intravenously at 10 μL/h beginning immediately after delivery of all pups, and the infusion was terminated 24 hours later by aspiration of the remaining reservoir contents. This protocol ensured a standardized 24-hour postpartum exposure window regardless of individual variability in delivery timing. Detailed pump programming parameters and the rationale for choice of OT dose are provided in Supplementary Methods and **Fig. S1**.

### 2. Impact of PPH-OT on maternal care and breast OT receptor (OTR) signaling

Maternal nurturing behavior was evaluated with the pup retrieval test, licking and grooming assay, and the weigh-suckle-weigh assay to assess lactational efficiency. We also quantified pup weight gain trajectories from birth to weaning as a surrogate measure of maternal care. Lastly, given that prolonged OT exposure can induce OT receptor (OTR) downregulation,^33^ we evaluated whether PPH-OT affected OTR expression within the mammary ductal epithelium. We performed this by aseptically isolating the 4th mammary fat pad immediately after weaning the pups, as described by Tovar et al.^34^ TaqMan^®^ qPCR analysis of OTR mRNA expression was performed using a custom probe as described by us previously.^30, 35^ Relative mRNA expression, normalized to the geometric mean of 5 reference genes (*Gapdh*, *Pgk1*, *PUM1*, *RPL13A*, and *Actb*), was calculated using the 2^-ΔΔCT^ method.

### 3. Breast milk transfer and quantification of OT

#### 3a. OT EIA assay in breastmilk

As a proof-of-principle experiment, we first assessed if systemically administered OT in a nursing postpartum (PP) dam would be recoverable in the breast milk. To investigate this, we intraperitoneally administered 50 I.U. OT or equivalent amount of saline in PP day 14 dams, when nursing routines were fully established, followed 1 hour later by collection of breast milk as described by Paul HA et al.^36^ Samples were assayed for OT using an enzyme immunoassay kit (K048-H1, Arbor Assays, Inc.) according to manufacturer’s instructions.

#### 3b. Stable isotope–labeled (SIL) OT infusion

Because ELISA-based OT cannot distinguish exogenously administered OT from the pulsatile endogenous OT released during lactation, we performed complementary experiments using stable isotope-labeled oxytocin (SIL-OT; [¹³C,¹⁵N]-Ile⁸-oxytocin; Biosynth International, Inc., Gardner, MA, USA), which is quantifiable by LC-MS/MS, allowing unequivocal identification and quantification of maternally administered OT independently of endogenous release. Both cysteine residues of SIL-OT were carbamidomethylated to prevent oxidative artifacts and ensure stability during LC-MS/MS analysis. Briefly, we implanted iPrecio^®^ SMP-200 microprocessor-controlled infusion pumps at GD16 and connected it to the right internal jugular vein under anesthesia as previously described.^30–32^ Immediately after delivery of all the pups at GD22, we aspirated the heparinized saline from the pump reservoir and replaced it with SIL-OT (1.6 mg/mL), which was subsequently infused at the preset rate of 10μL/h for 24h. Breast milk, approximately 0.8 to 1.25 mL/dam, was collected from the lactating dams at the completion of 24h of SIL-OT infusion. The collected milk fractions were stored at −80°C until further processing and LC-MS/MS analysis. Simultaneously, we collected maternal blood and processed the plasma for SIL-OT as previously described by us.^30^

#### 3c. LC-MS/MS assay for SIL-OT

To evaluate the presence of SIL-OT in breast milk, samples were processed using a specialized extraction protocol involving skimming, protein precipitation, and solid-phase extraction (SPE). Plasma samples were processed as described by us previously.^30^ LC-MS/MS analyses were performed on a Vanquish Neo nano UHPLC coupled to an Exploris 480 Orbitrap (Thermo Scientific, Waltham, MA, USA). Quantification was performed by LC–MS/MS using SIL-OT and a carbamidomethylated OT internal standard (CAM-OT), based on peak area ratios with defined quantifier and qualifier transitions. Details of the assay are provided in Supplementary Methods.

### 4. OTR expression analysis in newborn cortex

Based on our previous work demonstrating OTR downregulation in the developing brain after OT during labor induction,^30^ we investigated whether postnatal OT exposure produces a similar effect. To examine this, we harvested whole cortical hemispheres from P2 pups for TaqMan® qPCR analysis of OTR mRNA expression after 24 h postpartum maternal infusion of either OT or saline. Processing of samples for OTR qPCR was done with a custom TaqMan® OTR probe as described by us previously.^30, 35^ After collection of these samples, litter size was standardized to 8 pups/dam with equal sex distribution for behavioral studies.

### 5. Offspring neurobehavioral assessment

Rat pups were assessed for neurobehavioral consequences of postpartum OT exposure at P8 and again through juvenility and adolescence. Male and female offspring from OT-exposed and unexposed dams were assessed for early communicative behavior with maternal isolation-induced ultrasonic vocalizations (USV), anxiety-like behaviors with elevated plus maze, social behaviors with social approach and social play, cognitive inflexibility with spontaneous alternations, sensorimotor gating with prepulse inhibition, and empathy-like behaviors with observational fear learning as described by us previously (**Supplementary Table S2**).^30^

### 6. RNA sequencing to assess transcriptomic changes in the offspring brain

We collected the mPFC and the hypothalamus from male and female littermates from PPH-OT and control groups immediately after weaning. The mPFC was isolated as described previously,^30^ while dissection of the hypothalamus was based on anatomically defined landmarks.^37^ RNA-seq analysis was performed as described by us previously,^30^ and elaborated in greater detail in the Supplementary Methods. RNA-seq reads were then aligned to the Ensembl mRatBN7.2.113 primary assembly with STAR version 2.7.11b. All sequencing data are available through the NCBI Gene Expression Omnibus (GEO) under accession number GSE319898.

### 7. Statistical analysis

Data outliers were detected and eliminated using ROUT (robust regression and outlier analysis) with Q set to 1%. For experiments where sex was not included as a variable, data were analyzed with either Welch’s t-test or Mann-Whitney test as appropriate. Behavioral data were analyzed as previously described by us (detailed in Supplementary Methods).^30^ The critical alpha value for all behavioral analyses was p ≤ 0.05 unless otherwise stated. Quantitative data were analyzed with Prism 11 for macOS (GraphPad Software Inc., San Diego, CA, USA) and behavioral data were analyzed with IBM SPSS Statistics (v.28, Chicago, IL, USA) and presented as mean ± SEM.

Differential expression of genes (DEG) was analysed using limma-voom (R/Bioconductor).^30^ Three contrasts were fitted per brain region within a single linear model: a collapsed main-effect contrast (OT vs. control); within-sex contrasts (OT vs. control in males and females separately); and a sex-by-treatment interaction contrast [(male oxytocin − male control) − (female oxytocin − female control)]. Differentially expressed genes were defined by FDR ≤ 0.05 and absolute linear fold change ≥ 1.3, retaining genes with biologically meaningful effect sizes while excluding statistically significant but negligibly small expression changes. Pathway enrichment was assessed using GAGE, which tests whether log2 fold changes within a gene set are directionally shifted relative to all expressed genes, independently of DEG thresholds. GO and KEGG databases were used as references. Pathway enrichment was performed on the collapsed and within-sex contrasts for both regions and not on the interaction contrast. To determine whether suppression of neurodevelopmental disorder (NDD)–associated genes reflected directional regulation rather than nonspecific transcriptional perturbation, we tested for enrichment of NDD risk genes among downregulated versus upregulated mPFC DEGs using one-tailed Fisher’s exact tests. Three curated databases were queried: (i) SFARI Gene (January 2026 release; scores 1-3 analyzed both pooled and by individual tier), (ii) Developmental Brain Disorders database (DBD; autism and ID/DD categories), and (iii) SysNDD (Definitive tier, with the autosomal dominant subset analysed separately). Enrichment analyses were not performed for the hypothalamus, as no canonical NDD risk genes were identified among DEGs in any contrast or direction (overlap n = 0 across all databases and significance thresholds). P-values were adjusted with the Benjamini–Hochberg method and pathway enrichment results were filtered at FDR ≤ 0.05. The NDD enrichment analysis script and reference database files used in this study are publicly available at https://github.com/Arvindpalanisamy/PPH-OT-NDD-Enrichment/blob/main/ndd_enrichment_analysis.py (commit: 912b10b).

## RESULTS

### Maternal care behavior and neonatal wellbeing

Because peripartum exposure to OT can influence maternal mood and behavior,^38–41^ we examined maternal nurturing behaviors as described by us previously (**Fig. 1A-I**).^30^ Litter sizes were comparable between PPH-OT vs. saline groups and there were no differences in the survival rates of pups (**Fig. 1B-C**). When maternal care was examined in more detail, dams in both groups retrieved their pups with comparable latency and efficiency (**Fig. 1D**). One PPH-OT dam that did not retrieve any pups was excluded from all subsequent maternal and offspring analyses. There were no differences in maternal licking and grooming scores (**Fig. 1E**), suggesting that PPH-OT dams were able to nurture their offspring like control dams. These behavioral observations demonstrating good maternal care were supported by the weigh–suckle–weigh assay, which showed that lactational efficiency was not altered by PPH-OT exposure (**Fig. 1F**). Furthermore, exposure to PPH-OT did not downregulate OTR expression in the mammary tissue of the postpartum dam (**Fig. 1G**). OT-exposed pups showed a trend toward lower body weight during the first postnatal week that did not reach statistical significance (effect of treatment: F(1, 7) = 5.2, p = 0.057), followed by normalization before weaning (treatment × time interaction: (F(1.4, 9.5) = 0.19, p = 0.75; **Fig. 1H**). Guided by our prior observation that labor induction with OT reduces OTR gene expression in the newborn cortex,^30^ we examined whether postnatal OT exposure through breast milk would similarly impact the developing brain.

**Figure 1.**
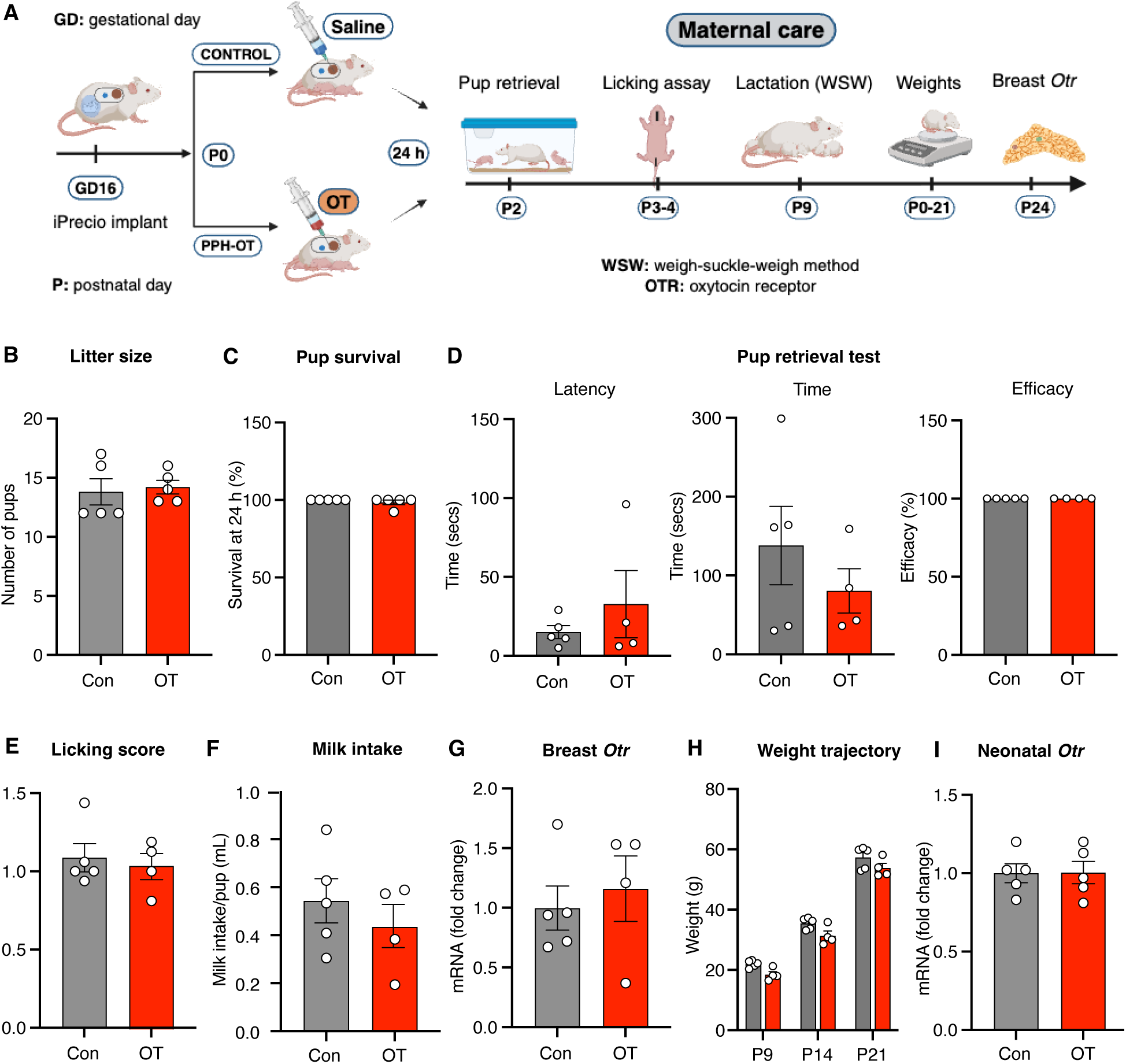
**Postpartum oxytocin exposure does not impair maternal care or wellbeing of the offspring**. **(A)** Experimental schematic illustrating peripartum oxytocin (PPH-OT) exposure via implanted pump and subsequent assessment of maternal behavior, lactation, and offspring outcomes. **(B–C)** Litter size and pup survival at 24 hours were comparable between control (saline) and PPH-OT groups (n = 5 dams per group), indicating no effect on early neonatal viability. **(D)** Maternal pup retrieval behavior was unchanged, with similar latency, total retrieval time, and retrieval efficiency between groups (n = 4-5 dams per group). **(E)** Licking and grooming behavior did not differ between control and PPH-OT dams (n = 4-5 dams per group). **(F)** Lactational performance assessed by the weigh–suckle–weigh (WSW) assay showed no differences in milk intake per pup (n = 4-5 dams per group). **(G)** Oxytocin receptor (OTR) expression in mammary tissue of postpartum dams was not altered by PPH-OT exposure (n = 4-5 dams per group). **(H)** Offspring exposed to PPH-OT exhibited reduced body weight during the first postnatal week but demonstrated appropriate catch-up growth by weaning. Mean pup weight increased significantly across postnatal development (P9–P21; two-way repeated-measures ANOVA, main effect of time: F(1.4, 9.5) = 1736, p < 0.0001), with no significant effect of treatment (F(1, 7) = 5.2, p = 0.0566) and no treatment × time interaction (F(1.4, 9.5) = 0.19, p = 0.75). **(I)** Neonatal cortical OTR gene expression was unchanged following PPH-OT exposure (n = 5 pups per group). Data are presented as mean ± SEM with individual data points.

In contrast to labor induction with OT, PPH-OT did not decrease OTR gene expression in the neonatal cortex (**Fig. 1I**), suggesting that the route, timing, or the magnitude of OT exposure may differentially affect neonatal cortical OTR signaling. Taken together, these findings indicate that PPH-OT did not impair maternal nurturing behavior but had a transient and reversible impact on neonatal wellbeing.

### Breast milk transfer of OT

We next asked whether maternally administered OT is secreted into the breast milk (**Fig. 2A-C**). We tested this in PP14 nursing dams and noted that 50 I.U. of systemically administered OT resulted in a significant increase in breast milk OT concentrations (**Fig. 2C**). To more directly track synthetic OT transfer, we performed complementary experiments using a 24-hour infusion of stable isotope–labeled oxytocin (SIL-OT) during the immediate postpartum period (**Fig. 2D-H**). Targeted LC-MS/MS analysis of breast milk samples allowed us to identify and quantify SIL-OT in 2 out of 3 treated samples, whereas all control samples were below the level of quantification. Despite inter-animal variability in measured concentrations, the reproducible detection of SIL-OT across treated animals confirmed that systemically administered OT is transferred into the mammary compartment. In a representative paired sample (**Fig. 2E**), concurrent plasma and milk measurements yielded a spot milk-to-plasma concentration ratio of approximately 0.71. Because samples were collected at a single, unsynchronized time point relative to nursing, this value does not reflect milk-to-plasma partition coefficient, which typically requires time-integrated pharmacokinetic analysis. Nonetheless, the observed ratio indicated that synthetic OT could reach milk at concentrations approaching those in the maternal circulation at the time of sampling, supporting physiologically meaningful transfer during lactation.

**Figure 2.**
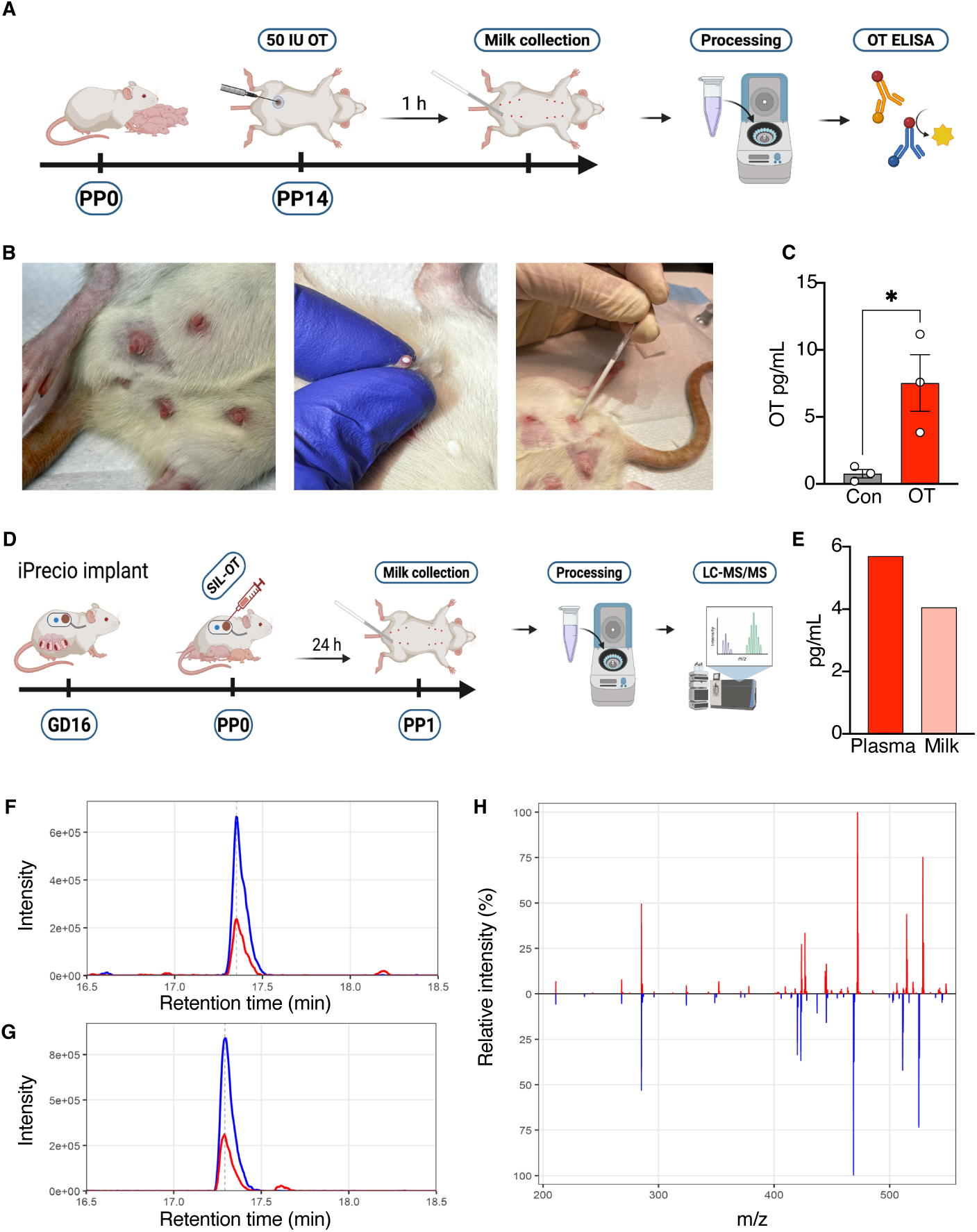
Lactational transfer of synthetic oxytocin confirmed by immunoassay and LC-MS/MS. **(A)** Experimental schematic for EIA-based OT detection. **(B)** Photographs illustrating the sequential steps of milk collection: mammary gland preparation, oxytocin-induced milk letdown and expression, and manual extraction with capillary tube. **(C)** Breast milk OT concentrations (pg/mL) in control (Con) and OT-treated dams at PP14. OT-treated dams showed significantly elevated milk OT levels compared to controls (*p = 0.03) (n=3 each). Data are mean ± SEM; individual data points shown. **(D)** Experimental schematic for SIL-OT quantification by LC-MS/MS. **(E)** SIL-OT concentrations (pg/mL) in a representative paired maternal plasma and breast milk sample collected at PP1, demonstrating lactational transfer of synthetic oxytocin. The spot milk-to-plasma ratio in this representative pair was approximately 0.71. **(F–H)** LC-MS/MS chromatograms and MS/MS spectral comparison of OT and SIL-OT. Extracted ion chromatograms (XIC) of OT (blue) and SIL-OT (red) are shown for **(F)** milk and plasma, demonstrating similar retention behavior with distinct signal intensities. **(H)** Mirror plot of representative MS/MS spectra for OT and SIL-OT showing closely matched fragmentation patterns and confirming analyte identity (n=2 control and 3 SIL-OT). PP: postpartum day.

### Neurobehavioral changes in the offspring

To determine whether postnatal oxytocin exposure (PPH-OT) alters neurobehavioral development, male and female offspring were evaluated across a developmental battery spanning early-life communication, juvenile social behavior, and adult behavioral domains (**Fig. 3A-O**). During maternal isolation at P8, OT-exposed pups emitted a greater number of ultrasonic vocalizations (USVs). However, this increase was attributable to reduced body temperature and was no longer significant after temperature adjustment (**Fig. 3B and Fig. S2**), indicating that this was physiologically mediated. The most prominent behavioral effects emerged during juvenile dyadic interactions (**Fig. 3F–L and Fig. S3**). OT-exposed offspring produced fewer total USVs (p = 0.05), accompanied by marked alterations in acoustic structure, including reduced average call frequency (p = 0.006), shorter call duration (p = 0.03) and phrase duration (p = 0.04), and increased frequency bandwidth (p = 0.0002). Call-type composition was also shifted toward a more aversive profile. OT-exposed offspring generated a higher proportion of aversive 22-kHz calls (p = 0.01), an effect that was more pronounced in females (p = 0.009), alongside a reduction in 50-kHz affiliative calls (combined p = 0.003; female p = 0.02). These findings indicate a selective disruption of social communication characterized by reduced call output, altered spectral features, and a shift toward negatively valenced signaling. In the social approach task (**Fig. 3N**), both groups retained a preference for the social over the empty chamber (**Fig. S4**). However, PPH-OT produced a significant treatment-by-sex interaction (p = 0.04), driven by reduced social preference in males (p = 0.01), indicating a sex-specific attenuation of sociability despite preserved overall social approach. In contrast, social play behavior (**Fig. 3D and Fig. S5-7**) was strongly sex-dependent but unaffected by treatment.

**Figure 3.**
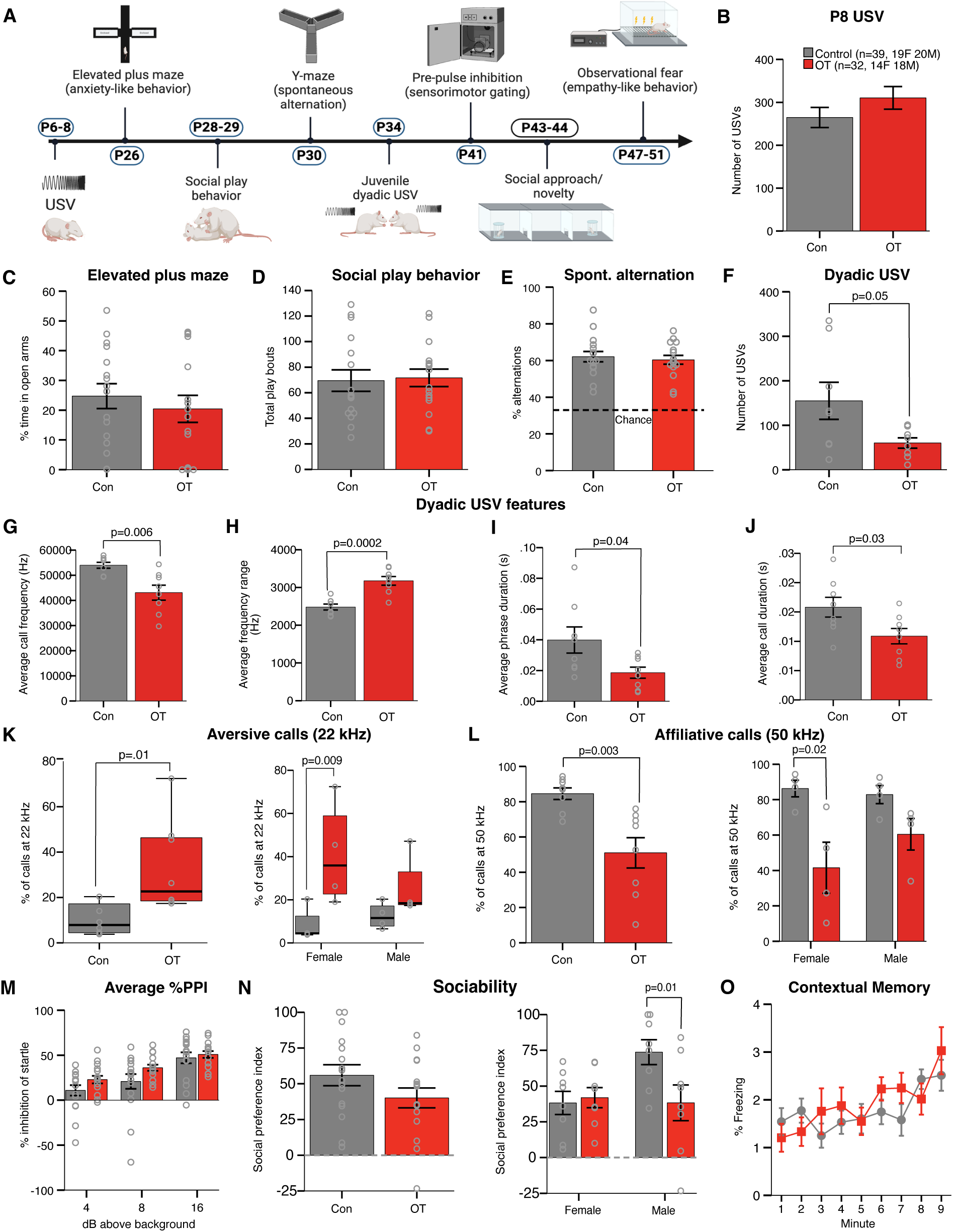
Postnatal oxytocin exposure selectively alters social communication and sociability. **(A)** Schematic of the experimental timeline illustrating behavioral assays across development, including early-life ultrasonic vocalizations (USVs), juvenile social play and dyadic communication, and adult behavioral testing. **(B)** At postnatal day 8 (P8), pups exposed to PPH-OT emitted a greater number of USVs during maternal isolation; this increase was attributable to reduced body temperature and was no longer significant when temperature was included as a covariate. **(C)** Anxiety-like behavior assessed by elevated plus maze (% time in open arms) was not different between groups. **(D)** Social play behavior (total play bouts) showed robust sex differences but was not altered by PPH-OT exposure. **(E)** Spontaneous alternation in the Y-maze was unchanged, indicating intact working memory. **(F)** During juvenile dyadic interactions, OT-exposed offspring emitted fewer total USVs (p = 0.05). **(G–J)** Acoustic analysis of dyadic USVs revealed reduced average call frequency (p = 0.006), increased frequency range (p = 0.0002), shorter phrase duration (p = 0.04), and shorter call duration (p = 0.03) in OT-exposed animals. **(K–L)** Call-type composition was shifted, with an increased proportion of aversive 22-kHz calls (combined p = 0.01; female p = 0.009) and a reduction in affiliative 50-kHz calls (combined p = 0.003; female p = 0.02). **(M)** Sensorimotor gating assessed by prepulse inhibition (PPI) was not affected by treatment. **(N)** In the social approach assay, both groups exhibited a preference for the social chamber; however, a treatment-by-sex interaction (p = 0.04) revealed reduced social preference in OT-exposed males (p = 0.01). **(O)** Contextual fear memory (% freezing) was not altered by PPH-OT exposure. Data are presented as mean ± SEM with individual data points overlaid. Statistical comparisons were performed as described in Methods, with p-values indicated in panels. Sample sizes: P8 USV (Control n = 39, OT n = 32), juvenile dyadic USV (Con = 8 pairs, OT n = 8 pairs), and all other behavioral assays (Con n = 16, OT n = 16), with all pups derived from 5 control and 4 PPH-OT exposed dams.

Males exhibited higher levels of play initiation, pouncing, and total play bouts compared to females, and these differences were preserved following OT exposure, with no effect on play metrics or reciprocity. Across multiple additional domains, including anxiety-like behavior (**Fig. 3C and Fig. S8**; elevated plus maze), spontaneous alternation (**Fig. 3E**; working memory), sensorimotor gating (**Fig. 3M and Fig. S9**; prepulse inhibition), and contextual fear memory (**Fig. 3O and Fig. S10**), no significant effects of PPH-OT were observed. Collectively, these data demonstrate that postnatal oxytocin exposure produces selective, domain-specific alterations in social communication and sociability, with effects that are context-dependent and sexually dimorphic, while sparing core cognitive, affective, and sensorimotor function.

### Transcriptomic changes in the mPFC

Given the observed alterations in social behavior, we examined transcriptional changes in the mPFC, a key hub for social processing (**Fig. 4**).^42^ PPH-OT induced widespread transcriptional changes (**Fig. 4A**), with 3,519 differentially expressed genes (DEGs; 2,234 upregulated and 1,285 downregulated). Sex-stratified analysis revealed a marked dimorphism in response magnitude (**Fig. 4B**). Males exhibited 1,732 DEGs (1,372 upregulated and 360 downregulated), whereas females showed 693 DEGs (298 upregulated and 395 downregulated), representing an approximately 2.5-fold difference. Beyond scale, the transcriptional responses were qualitatively distinct (**Fig. 4C**). Only 132 DEGs were shared between sexes, indicating largely non-overlapping transcriptional programs. Consistent with this divergence, sex-by-treatment interaction analysis identified 1,229 genes with significantly sex-differentiated responses (847 male-biased and 382 female-biased). Pathway analysis further underscored these differences (**Fig. 4D**). In males, 258 GO biological process terms were significantly downregulated, dominated by pathways related to nervous system development, synapse organization, neurogenesis, axon guidance, dendritic spine organization, and transcriptional regulation. In addition, the endoplasmic reticulum (ER) protein processing pathway was selectively suppressed in males, with all nine unfolded protein response (UPR) gene hits absent from the female DEG set. In contrast, the female transcriptional response was characterized by suppression of cilia- and microtubule-based processes and enrichment of immune system pathways and extracellular matrix organization, a profile not observed in males. All mPFC-related data files are included in the Supplementary File (**Data S1-S13**).

**Figure 4.**
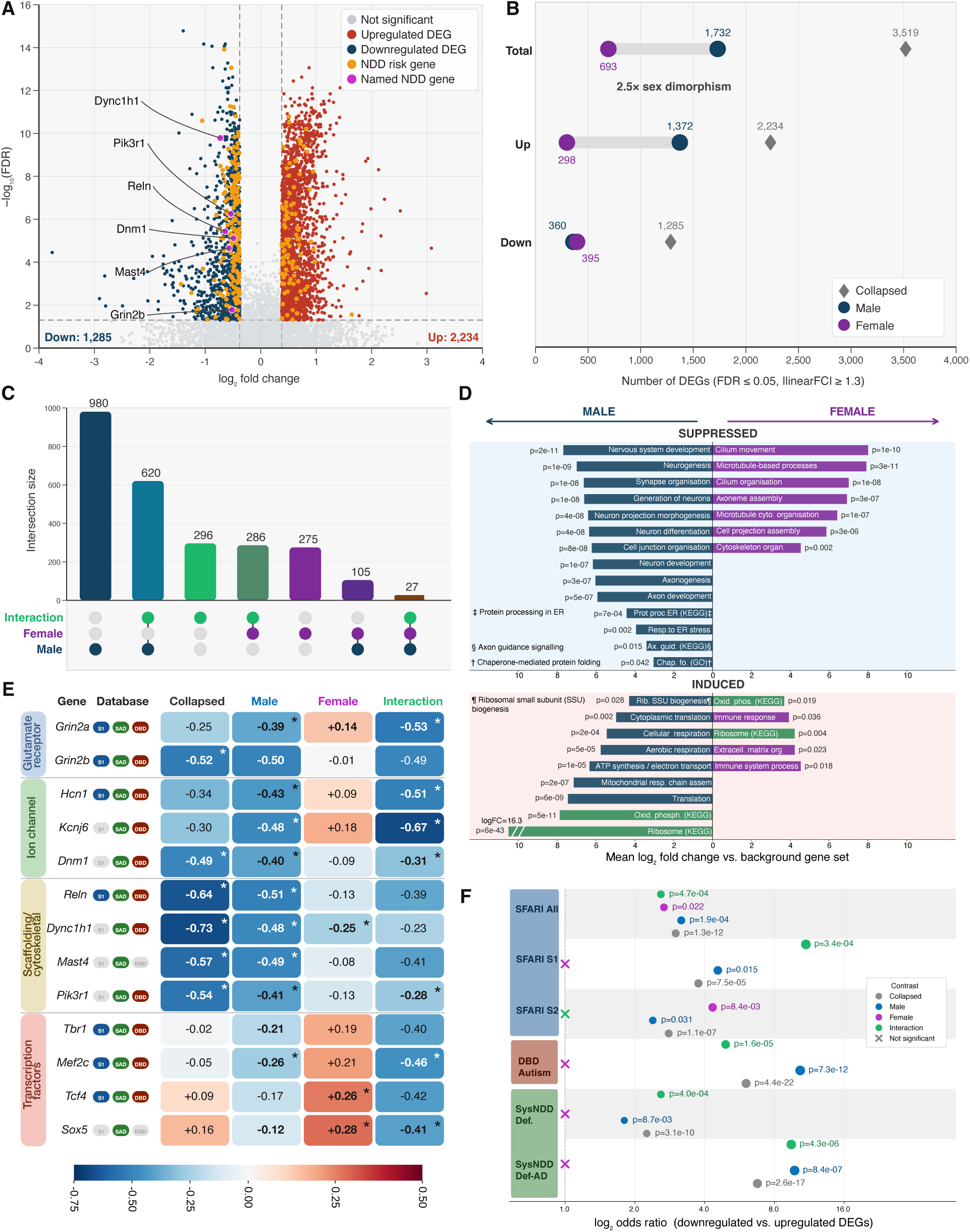
Postpartum oxytocin exposure induces sex-specific transcriptional and neurodevelopmental alterations in the medial prefrontal cortex. **(A)** Volcano plot of differentially expressed genes (DEGs) in the medial prefrontal cortex (mPFC) following postpartum oxytocin exposure (PPH-OT) compared with controls (n=10 pups per sex per condition, with pups derived from at least 9 independent litters). A total of 3,519 genes were differentially expressed (2,234 upregulated and 1,285 downregulated), with selected neurodevelopmental disorder (NDD)–associated genes highlighted. **(B)** Sex-stratified DEG counts showing a greater magnitude of transcriptional change in males (1,732 DEGs; 1,372 upregulated and 360 downregulated) compared with females (693 DEGs; 298 upregulated and 395 downregulated). **(C)** UpSet plot illustrating overlap of DEGs across male, female, and interaction contrasts. Limited overlap is observed, with 132 shared genes and 1,229 genes exhibiting significant sex-by-treatment interaction. **(D)** Gene ontology (GO) and pathway enrichment analysis of DEGs in males (left) and females (right). **(E)** Heatmap of selected NDD-associated genes showing differential expression (log₂ fold change) across collapsed, male, female, and interaction contrasts. Genes are grouped by functional category, including glutamatergic signaling, ion channels, cytoskeletal/scaffolding proteins, and transcriptional regulators. Database annotations (SFARI score-1, SysNDD autosomal dominant, DBD autism) are indicated, with genes not in curated databases shown in gray. Asterisks denote FDR ≤ 0.05 and |linear fold change| ≥ 1.3. **(F)** Forest plot showing enrichment of NDD-associated gene sets among downregulated DEGs (one-tailed Fisher’s exact test, FDR ≤ 0.05). In the collapsed analysis, downregulated genes are enriched for SFARI autism risk genes (OR = 3.00, p = 1.4×10⁻¹²), DBD autism genes (OR = 6.04, p = 4.4×10⁻²²), and SysNDD definitive genes (OR = 2.25, p = 3.1×10⁻¹⁰), with the strongest enrichment observed for SysNDD autosomal dominant genes (OR = 6.76, p = 2.6×10⁻¹⁷). In contrast, female downregulated genes show minimal enrichment across most NDD-gene sets, with odds ratios approximating 1 (pink crossed markers).

To determine whether these transcriptional changes preferentially involved disease-relevant genes, we assessed enrichment of neurodevelopmental disorder (NDD) gene sets among downregulated DEGs (**Fig. 4F** and **Data S14**). In the collapsed analysis, downregulated genes were enriched for SFARI autism risk genes (OR = 3.00, p = 1.4×10⁻¹²), DBD autism genes (OR = 6.04, p = 4.4×10⁻²²), and SysNDD definitive genes (OR = 2.25, p = 3.1×10⁻¹⁰), with the strongest enrichment observed for SysNDD autosomal dominant genes (OR = 6.76, p = 2.6×10⁻¹⁷). This enrichment was driven predominantly by males. Male downregulated genes were significantly enriched across all databases and confidence tiers, including SFARI all, SFARI score-1, DBD autism, and SysNDD autosomal dominant genes, whereas female downregulated genes showed no enrichment across high-confidence NDD categories (all p > 0.05), with only modest enrichment observed for broader SFARI categories. Gene-level analysis confirmed this pattern (**Fig. 4E**). Male downregulated genes included multiple high-confidence NDD risk genes, including *Grin2a*, *Grin2b*, *Hcn1*, *Dnm1*, *Kcnj6*, *Reln*, *Mast4*, *Pik3r1*, and *Dync1h1*, with additional transcriptional regulators (*Tbr1*, *Mef2c*, *Tcf4*, and *Sox5*) identified through the interaction contrast. These genes span key functional categories, including glutamatergic signaling, ion channels, cytoskeletal organization, and transcriptional regulation. Collectively, these results define a sex-specific transcriptional response to PPH-OT in the mPFC, characterized by selective suppression of neurodevelopmental gene programs and enrichment of NDD-associated genes in males.

### Transcriptomic changes in the hypothalamus

Given the central role of the hypothalamus in OT-OTR signaling,^43^ we evaluated transcriptional changes in this region (**Fig. 5**). Differential expression analysis (**Fig. 5A**) identified 402 hypothalamic DEGs (108 upregulated, 294 downregulated), indicating a predominantly suppressive transcriptional response. Distinct features included a coordinated catecholaminergic upregulation module (*Slc6a3*, *Th*, *Ddc*, *Slc18a2*, *En1*, *Chrna5/6*) and a concomitant suppression of unfolded protein response (UPR) genes (*Hspa5*, *Hspa1a*, *Pdia4*, *Creld2*), with *Npw* representing the principal shared upregulated gene across sexes. The hypothalamic response exhibited marked sexual dimorphism, with 141 DEGs in males (31 up, 110 down) compared to only 12 in females (4 up, 8 down), a 11.8-fold difference in magnitude (**Fig. 5B**). Cross-sex comparison (**Fig. 5C-D**) revealed that the transcriptional response was overwhelmingly male-driven, with 135 male-specific DEGs, 6 female-specific DEGs, and only 6 shared genes. Shared genes followed the line of concordance, whereas the catecholaminergic activation and UPR suppression signatures were largely confined to males, indicating sex-specific engagement of both neurotransmitter and proteostasis pathways. Pathway analysis (**Fig. 5E**) demonstrated a clear divergence in biological programs. In males, upregulated pathways were restricted to synaptic signaling and chemical synaptic transmission, with no detectable downregulated pathways. In contrast, females exhibited a coherent systems-level response, with enrichment of synaptic signaling, neurogenesis, nervous system development, and neurotransmitter receptor activity, along with cholinergic synapse signaling. Notably, translational pathways were oppositely regulated across regions and sexes: ribosomal programs were suppressed in females, contrasting with the activation observed in the male mPFC, suggesting region- and sex-specific reprogramming of protein synthesis. The sex-by-treatment interaction identified only three genes (*Sdf2l1*, *Serpinh1*, *Bcat2*), all reduced in males (**Fig. 5F**). *Sdf2l1* and *Serpinh1* overlapped with the mPFC interaction signature, identifying a conserved cross-regional axis of male-biased UPR suppression. In contrast to the mPFC, hypothalamic DEGs showed no enrichment for neurodevelopmental disorder risk genes across multiple curated databases, indicating that transcriptional vulnerability is region-specific. All hypothalamus-related data files are included in the Supplementary File (**Data S15-S27**).

**Figure 5.**
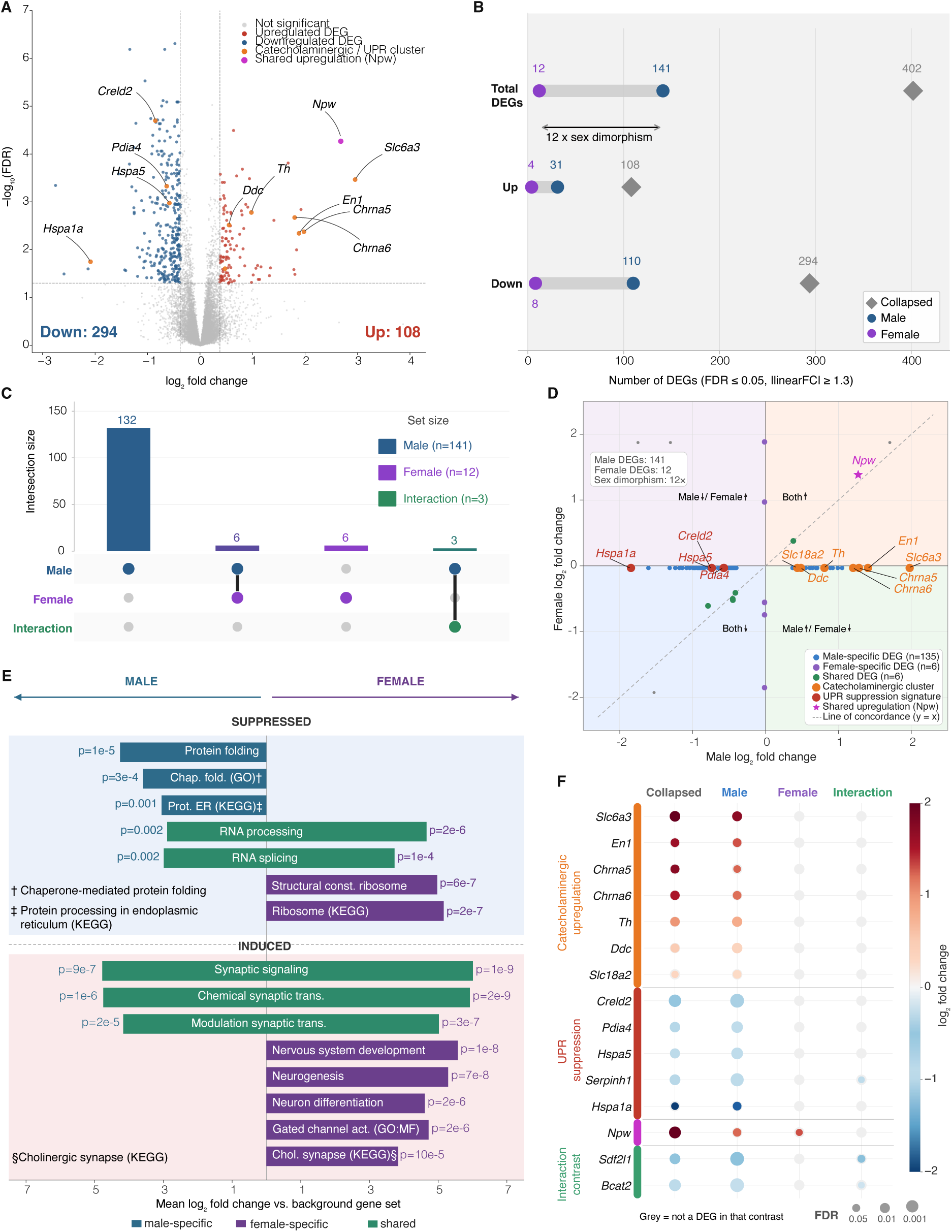
Postpartum oxytocin exposure induces sex-specific transcriptional reorganization of the hypothalamus. **(A)** Volcano plot of differentially expressed genes (DEGs) in the hypothalamus following PPH-OT compared with sex-matched controls (n=10 pups per sex per condition, with pups derived from at least 9 independent litters). A total of 402 genes were differentially expressed (108 upregulated and 294 downregulated). Two functionally distinct clusters are highlighted: a catecholaminergic/UPR cluster (orange) comprising genes involved in dopamine neurotransmission and endoplasmic reticulum stress, and Npw (magenta), the sole gene upregulated in both sexes. **(B)** Sex-stratified DEG counts illustrating a 12-fold sex dimorphism: males showed 141 DEGs (31 upregulated and 110 downregulated) compared with 12 DEGs in females (4 upregulated and 8 downregulated), with the collapsed contrast yielding 402 DEGs (108 upregulated and 294 downregulated). **(C)** UpSet plot showing overlap of DEGs across male, female, and interaction contrasts. The male-specific set accounts for the dominant signal (132 genes), with limited overlap across contrasts (6 male–female shared, 6 male–interaction shared, 3 in all three). **(D)** Sex-dimorphic DEG landscape plotted as male versus female log₂ fold change for each gene. Male-specific DEGs (n = 135, blue) cluster along the x-axis, reflecting absent female responses; female-specific DEGs (n = 6, purple) are distributed along the y-axis; shared DEGs (n = 6, grey) show concordant regulation in both sexes. The catecholaminergic gene cluster (orange; *Th*, *Ddc*, *Slc6a3*, *Slc18a2*, *En1*, *Chrna5*, *Chrna6*) is selectively induced in males, and a UPR suppression signature (red; *Hspa5*, *Hspa1a*, *Creld2*, *Pdia4*) is selectively suppressed in males. *Npw* (magenta star) was the sole gene upregulated in both sexes. The dashed diagonal denotes the line of concordance (y = x). **(E)** Butterfly bar chart of GO-BP and KEGG pathway enrichment in males (left) and females (right) for suppressed (top, blue background) and induced (bottom, pink background) DEG sets. Shared pathways are shown in green bars. Bar length reflects mean log₂ fold change relative to the background gene set; p-values are shown at bar tips. **(F)** Dot plot of named gene clusters showing log₂ fold change across collapsed, male, female, and interaction contrasts. Genes are grouped by functional cluster: catecholaminergic upregulation, UPR suppression, shared upregulation (*Npw*), and interaction contrast (*Sdf2l1*, *Bcat2*). Dot color indicates log₂ fold change and dot size reflects FDR (large, FDR ≤ 0.001; medium, FDR ≤ 0.01; small, FDR ≤ 0.05). Grey dots indicate that the genes did not reach DEG threshold in that contrast.

### Divergent regional transcriptomic responses to PPH-OT exposure

Having characterized transcriptional changes in the mPFC and hypothalamus independently, we next performed a direct cross-regional comparison (**Fig. 6**). The magnitude of transcriptional response differed markedly between regions, with 3,519 DEGs in the mPFC compared to 402 in the hypothalamus (**Fig. 6A**). These responses also showed opposing directionality: mPFC changes were activation-dominant (63% upregulated), whereas hypothalamic responses were largely suppressive (73% downregulated), particularly in males (**Fig. 6B**). Sex-stratified analysis revealed region-specific dimorphism. In the mPFC, males exhibited a 2.5-fold greater DEG burden than females (1,732 vs. 693 DEGs; 71% male-biased), whereas the hypothalamus showed a more pronounced 11.8-fold male bias (141 vs. 12 DEGs; 92% male-biased) (**Fig. 6C**). Pathway analysis further distinguished the two regions. In the mPFC, male downregulated genes were enriched for neurodevelopmental and synaptic processes, while upregulated pathways included ribosomal and translational programs. In contrast, hypothalamic responses were more restricted and characterized by male-specific suppression of proteostasis pathways (UPR/ER chaperone signaling) alongside induction of catecholaminergic and synaptic signaling pathways. Female hypothalamic responses were limited but showed coherent pathway-level changes that were often distinct from those observed in the mPFC (**Fig. 6D**). Consistent with these differences, downregulated mPFC DEGs were significantly enriched for neurodevelopmental disorder (NDD) risk genes across multiple databases, whereas hypothalamic DEGs showed no enrichment (**Fig. 6E**). Overlap between regions was limited, with 130 shared genes (32% of hypothalamic DEGs; ∼4% of mPFC DEGs) (**Fig. 6F**). However, shared genes exhibited moderate concordance in directionality (r = 0.65; 79% concordant), indicating partial alignment despite divergent global programs (**Fig. 6G**). Together, these findings demonstrate that PPH-OT exposure induces broad, sex-biased, NDD-relevant transcriptional remodeling in the mPFC, whereas the hypothalamic response is comparatively constrained, male-dominated, and centered on proteostasis and catecholaminergic signaling.

**Figure 6.**
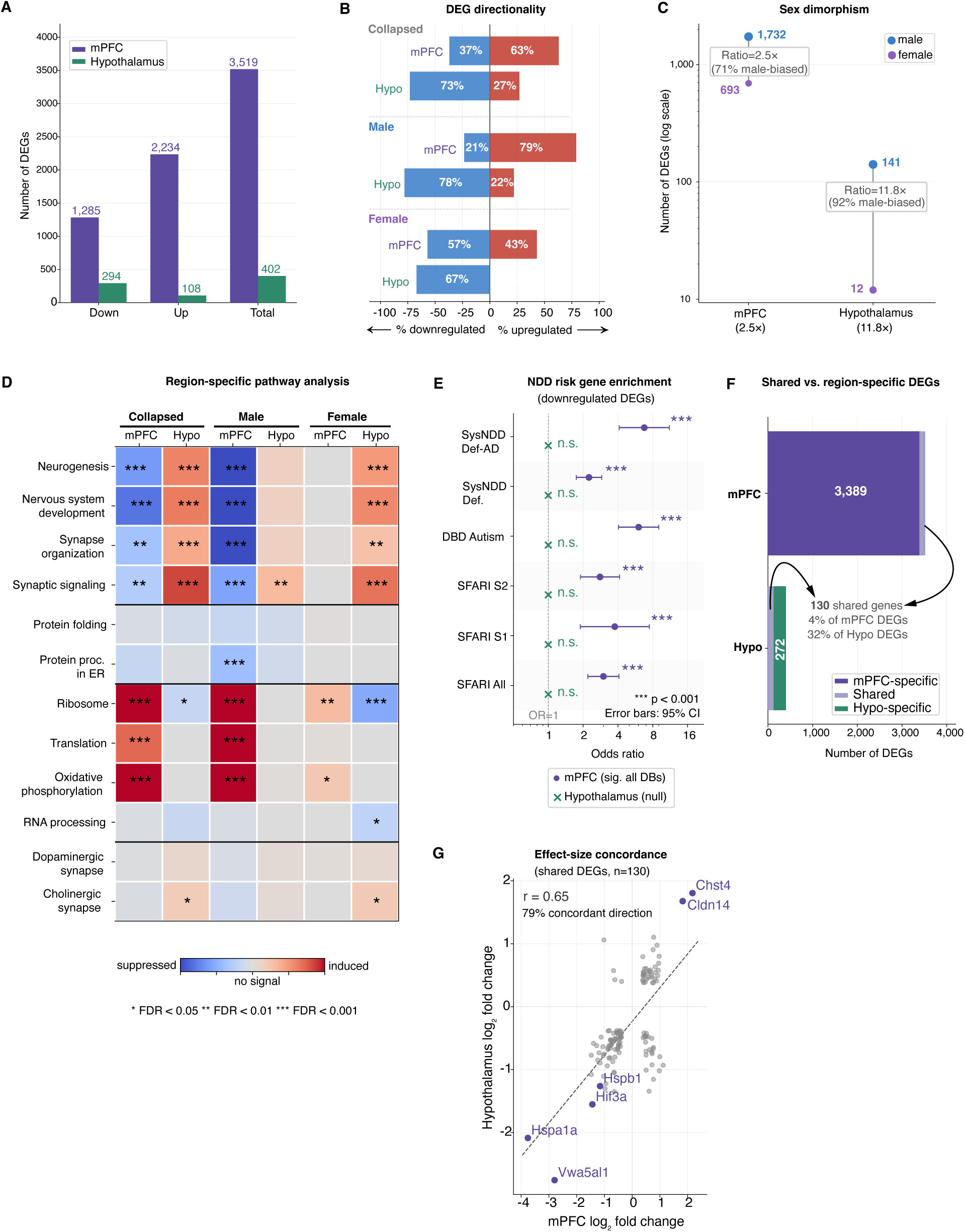
Divergent regional transcriptomic responses to postnatal oxytocin exposure. **(A)** DEG burden in mPFC (3,519 genes) and hypothalamus (402 genes) under the collapsed contrast. **(B)** Directionality of transcriptional change shows activation-dominant responses in mPFC (63% upregulated) and suppression-dominant responses in hypothalamus (73% downregulated). **(C)** Sex-stratified analysis reveals greater male bias in hypothalamus (11.8x) than mPFC (2.5x). **(D)** Pathway analysis (GAGE) identifies region- and sex-specific programs, with mPFC enriched for neurodevelopmental and synaptic processes and hypothalamus showing proteostasis-related suppression and catecholaminergic/synaptic induction. **(E)** Neurodevelopmental disorder (NDD) risk genes are enriched among downregulated mPFC DEGs across multiple databases, but not in hypothalamus (n.s.); error bars indicate 95% confidence intervals. **(F)** Limited overlap between regions (130 shared genes; 32% of hypothalamic DEGs, 4% of mPFC DEGs). **(G)** Shared genes show moderate effect-size concordance (r = 0.65; 79% concordant direction).

## DISCUSSION

Our preclinical study shows that postpartum exposure to OT in the context of PPH prophylaxis can have sex-specific neurobehavioral consequences for the developing brain despite preserved maternal behaviors, and demonstrates for the first time that exogenously administered PPH-OT can be lactationally transferred to the neonate. Furthermore, our data identify a sex- and brain region-specific transcriptional response to PPH-OT in the offspring, with the strongest effects localized to the male mPFC, where developmental gene programs governing neuronal maturation and circuit organization were selectively suppressed.

The major neurobehavioral impacts of PPH-OT on the offspring were two-fold: (1) altered social communication, and (2) male-specific changes in social approach. Rat pups typically generate two types of USVs: a 50 kHz affiliative call that facilitates social interactions and a 22 kHz aversive call that reflects pain, anxiety, or danger.^44–48^ A unique neurobehavioral feature we observed was the alteration in USVs when OT-exposed pups engaged in social play behavior with other OT-exposed pups. OT-exposed pups not only vocalized less, but generated a disproportionately higher number of 22 kHz distress calls, especially females. Another standout feature was the male-specific decrease in social preference in OT-exposed pups. Taken together with the lack of changes in other behavioral domains, our findings suggest that PPH-OT selectively and sex-specifically affects sociality and social communication in the offspring. Of note, these behavioral findings were distinct from our previous study examining intrapartum exposure to OT, when sociality was unaltered and empathy was the most affected neurobehavioral domain,^30^ suggesting that the route, magnitude, and chronological timing of exposure could be consequential.

The mPFC emerged as the primary locus for sexually dimorphic transcriptional change after PPH-OT. This is notable because the mPFC matures over an extended perinatal window, making it particularly vulnerable to early life disruption and frequently implicated in neurodevelopmental disorders such as autism spectrum disorders.^49, 50^ The male-specific pattern of NDD risk gene suppression we report parallels one of the most replicated and least explained epidemiological observations: the approximately 2-to-4:1 male-to-female prevalence ratio in autism spectrum disorder and other NDDs.^1–3^ In alignment with the female protective effect,^51^ our findings suggest that a ubiquitous postnatal hormonal exposure produces a sexually dimorphic transcriptional response in the developing mPFC that selectively suppresses NDD risk gene networks in males, while leaving females largely unaffected. Critically, this male-specific enrichment converged across three independent reference sets (SFARI, DBD, and SysNDD) and across three independent statistical contrasts. The particularly strong enrichment (OR ≈ 10) for autosomal dominant NDD genes in male downregulated DEGs, including core regulators of cortical circuit assembly (*Grin2a*, *Grin2b*, *Reln*, *Tbr1*, *Sox5*), merits special consideration. Dominant NDD mutations are high-penetrance by definition and haploinsufficiency of a single allele is sufficient to disrupt neurodevelopment. Therefore, even partial reductions in expression during a critical developmental window could have phenotypic consequences. We do not propose that PPH-OT causes autism. Rather, we identify oxytocinergic modulation of prefrontal NDD risk gene expression as a sexually dimorphic process that operates in a direction consistent with the known sex bias in NDD prevalence.

Given the established influence of maternal nurturing on offspring neurobehavioral development,^52^ and the ongoing debate regarding the effects of exogenous OT on breastfeeding outcomes,^53–55^ maternal care behavior was robustly assessed as a potential confound. Detailed maternal behavioral phenotyping and preserved OTR expression in mammary ductal epithelium together indicated that PPH-OT did not disrupt maternal nurturing. Therefore, we examined the next logical possibility that OT could be transferred through breast milk, a notion supported by the presence of radioactive OT in the gastric content of pups after intraperitoneal administration in postpartum dams.^22^ In our study, we intravenously administered and directly confirmed the lactational transfer of SIL-OT for the first time. This strongly supports the possibility that the increase in plasma OT in pups after maternal OT reflects absorption from the neonatal gastrointestinal tract, presumably through the receptor for advanced glycation end-products (RAGE) as shown by Higashida et al.^23^ Our work, therefore, highlights the need to better define neonatal OT exposure following postpartum administration.

In the absence of objective evidence for impaired maternal care, and given the trend toward lower body weight in OT-exposed pups, we speculate that transient neonatal weight suppression, if confirmed with adequate power, may reflect a direct effect of lactational OT exposure. One possible explanation is disruption of brown adipose tissue (BAT) thermogenesis, as OT is critical for BAT activation and maintenance of body temperature.^56–58^ Disruption of OT signaling, either through OT or OTR knockout,^59–61^ is associated with hypothermia. We speculate that prolonged neonatal OT exposure through lactation may transiently alter thermogenic regulation, a hypothesis that needs independent testing. This is not only supported by evidence for breast milk transfer of OT, but also by our observation that the decreased weight of OT-exposed pups is transient, with improvement and normalization before weaning.

Nevertheless, taken together with clinical studies suggesting impairment of neonatal feeding behaviors after perinatal OT exposure,^55, 62–65^ PPH-OT should be considered as a potential covariable when assessing breastfeeding outcomes.

Our study has a few limitations. First, rats do not experience PPH, and, therefore, our anthropomorphization must be interpreted with caution. We selected the lowest possible OT infusion dose that ensured authentic maternal nurturing behaviors without affecting milk production (cumulative dose ∼ 85 I.U.), which is at the higher end of clinical PPH-OT dosing ranges.^66, 67^ Second, subtle changes in maternal behaviors may have been missed by our assays. Cross-fostering could be a useful tool to delineate the impact of OT *per se*, but the act of cross-fostering in itself has been shown to alter neurodevelopmental and behavioral outcomes in the offspring.^68–70^ Third, although multiple analytical approaches converge on disrupted cortical maturation in male offspring, causal relationships between PPH-OT and specific cellular or circuit mechanisms remain to be established. Notably, a substantial body of evidence, including work from our laboratory, demonstrates pronounced sexual dimorphism within the OT–OTR system, which may contribute to sex-specific neural and behavioral responses to genetic or pharmacologic perturbations.^29, 30, 71–78^ Fourth, because our timed breast milk sample collection was not synchronized to nursing bout, the measured milk and plasma SIL-OT concentrations were spot values. A full milk-to-plasma ratio could not be calculated, as repeated timed milk expression was not feasible in lactating rats. Finally, we chose to examine the hypothalamus and the mPFC, considering the importance of these regions for OT-OTR biology,^43, 79–81^ and social behavior,^82–85^ respectively. PPH-OT–associated neurobehavioral changes might involve broader circuit-level alterations beyond these regions, warranting comprehensive 3D brain mapping coupled with high-resolution spatial transcriptomics analyses.

In summary, our study raises the concern that postpartum prophylaxis of hemorrhage with oxytocin may influence neurodevelopment through lactational transfer in a sex-dependent manner. These findings support the need for clinical studies to define synthetic oxytocin exposure in the nursing infant, determine milk-to-plasma ratios and the relative infant dose, evaluate the relative safety of high dose oxytocin use, and assess neurodevelopmental outcomes in children following such exposure.

## Conflict of Interest

The authors declare no conflict of interest.

## Ethics Approval

All animal experiments were approved by the Institutional Animal Care and Use Committee at Washington University in St. Louis (protocol #23-0021) and conducted in accordance with institutional and ARRIVE guidelines.

## Data Availability

All sequencing data are available through the NCBI Gene Expression Omnibus (GEO) under accession number GSE319898.The NDD enrichment analysis script and reference database files used in this study are publicly available at https://github.com/Arvindpalanisamy/PPH-OT-NDD-Enrichment/blob/main/ndd_enrichment_analysis.py (commit: 912b10b).

## Supplementary Information

Supplementary Information included as a separate file for availability at *Molecular Psychiatry*.

## Acknowledgments

This work was supported by the following NIH grant to AP (1R21HD117239-01A1). Animal behavioral studies were performed at the Model Systems Core of the Intellectual and Developmental Disabilities Research Center (IDDRC) at Washington University in St. Louis.

This facility is supported by the Eunice Kennedy Shriver National Institute Of Child Health & Human Development of the National Institutes of Health (P50 HD103525). RNA-seq was performed by the Genome Technology Access Center at McDonnell Genome Institute (GTAC@MGI), Washington University School of Medicine. The Center is partially supported by NCI Cancer Center Support Grant P30 CA91842 to the Siteman Cancer Center from the National Center for Research Resources (NCRR), a component of the National Institutes of Health (NIH), and NIH Roadmap for Medical Research. Mass Spectrometry analyses were performed by the Mass Spectrometry Technology Access Center at the McDonnell Genome Institute (MTAC@MGI) at Washington University School of Medicine, supported by the Diabetes Research Center/NIH grant P30 DK020579, Institute of Clinical and Translational Sciences/NCATS CTSA award UL1 TR002345, and Siteman Cancer Center/NCI CCSG grant P30 CA091842.

## Author contributions

A.P. conceived the study design, planned the experiments, and wrote the manuscript with help from all co-authors. G.S. and T.G. conducted the animal surgery, and performed the molecular biology experiments. M.L. and E.T. performed the statistical analysis and generated the figures and tables for RNA-seq data. Y.A.G., M.S., and R.W. performed and analyzed the mass spectrometry data from stable isotope labeled oxytocin experiments and helped draft the manuscript. K.B.M. conducted the behavioral experiments under the supervision of S.E.M.

S.E.M. analyzed the behavioral data, generated figures for the manuscript, and helped with interpretation and drafting of the manuscript. R.G. helped with literature review and writing the discussion. All authors actively participated in the drafting and editing of the manuscript.

## Declaration of generative AI and AI-assisted technologies during manuscript preparation

Enrichment analyses of neurodevelopmental disorder risk genes were performed using Python code written and executed with the assistance of Claude Sonnet 4.6 (Anthropic; March 2026), implementing Fisher’s exact test (scipy.stats.fisher_exact). All statistical outputs were verified by the authors. Manuscript drafting and editing were assisted by Claude Sonnet 4.6 and ChatGPT. AI tools were not used to generate or modify primary experimental data. The author(s) reviewed and edited the content as needed and take(s) full responsibility for the content of the published article. Schematic illustrations were created at https://BioRender.com.

## Notes

https://github.com/Arvindpalanisamy/PPH-OT-NDD-Enrichment/blob/main/ndd_enrichment_analysis.py

### Competing Interest Statement

The authors have declared no competing interest.

