## Supplementary material for "Lactational transfer of synthetic oxytocin and sex-specific suppression of neurodevelopmental risk gene networks in the rat prefrontal cortex": Supplementary Material.docx

**Supplementary Methods**

iPrecio programming

For iPrecio pump surgery and sham controls, both groups of animals were exposed to the same duration of anesthesia and received buprenorphine analgesia and antibiotics. In the PPH-OT group, the iPrecio pump was preprogrammed to infuse heparinized saline at 3 mL/h until midnight of GD21, and at 10 mL/h after to keep the pump tubing patent. Because of this infusion schedule, when saline was completely replaced with OT (600 mcg/mL) immediately after birth of the pups at GD22 (postnatal day P0), it was infused at a rate of 10 mL/h. We terminated the infusion by completely aspirating the OT from the pump reservoir 24 h after instillation. This approach allowed us to account for variability in the timing of birth among dams, and ensured that each dam received 24 h of intravenous OT infusion in the postpartum period at the pre-set rate. We performed additional experiments with 2x OT dose and noted a trend towards reduced lactational efficiency (**Fig. S1**). Therefore, the dose and duration of OT infusion that we chose was a compromise between clinical translatability and practicality, respectively.

**qPCR for breast OTR expression**

Briefly, the postpartum dam was deeply anesthetized with isoflurane and pinned to a Styrofoam board in the supine position. The 4th left nipple was identified counting down from the neck, and the skin around the nipple was incised and separated from the abdominal muscle carefully, keeping the mammary fat pad attached to the skin. The mammary fat pad was subsequently carefully microdissected to include the entire ductal tree from the proximal nipple region out to the distal branches and stored in -80°C. RNA was extracted using RNeasy Lipid Tissue Mini Kit (QIAGEN) and converted to cDNA using SuperScript IV VILO Master Mix kit (Invitrogen, Thermo Fisher Scientific) and 25 ng of the template cDNA was then combined with a ready-to-use TaqMan Fast Advanced qPCR Master Mix (Thermo Fisher Scientific) for gene expression experiments with a custom TaqMan® OTR probe. Thermal cycling was performed in 7500 Fast Real-Time PCR System (Applied Biosystems®) and the threshold cycle (Ct) values for all genes were calculated using proprietary software.

**Processing steps for breast milk OT ELISA**

We administered OT (2 I.U., i.p.) to the dam using a 25-gauge needle under brief isoflurane anesthesia to trigger milk let-down. Subsequently, we quickly prepared the area around the nipples (inguinal mammary glands 4-6 and 10-12) by shaving the fur. When milk let-down was visually confirmed, typically 10 min after i.p. OT injection, we expressed milk manually from the prepared nipples by gentle massage of the mammary glands while simultaneously collecting it using a capillary tube. Milk collected from each dam was pooled into a pre-chilled, low-protein-binding microcentrifuge tube until we obtained approximately 1mL/dam. The milk samples were centrifuged at 20,000 x g for 30 min at 4◦C to separate the creamy layer containing the fat on the top. The clarified milk was collected and transferred to a new microcentrifuge tube and stored at -80^°^C. In brief, each sample was diluted 1:10 in the assay buffer and applied to the designated well(s), respectively. All experiments were performed in duplicate, and the absorbance was read with Tecan Infinite M200 PRO (Tecan Group Ltd., Switzerland) at 450 nm. OT concentrations were determined by comparison with predetermined standards, corrected for the dilution factor and reported as pg/mL.

**LC-MS/MS quantification and validation**

Briefly, milk samples were thawed on ice and centrifuged at 5,000 × g for 10 min at 4 °C to remove the cream layer. The skim milk was transferred to new tubes and supplemented with EDTA (10 mM), NaCl (15 mM), and formic acid (0.1%), then spiked with 2 μL of a 0.1 μg/mL solution of carbamidomethylated unlabeled OT (CAM-OT) as the internal standard (IS). Protein precipitation was performed by adding 1 mL of acetonitrile, incubating at −20 °C for 1 hour, and centrifuging at 21,000 × g for 10 min. The supernatant was subjected to liquid-liquid extraction with MTBE twice, and the resulting hydrophilic phase was dried and reconstituted in 2 mL of 0.1% formic acid for cleanup. SPE was carried out using Oasis HLB cartridges (3 cc, 150 mg) conditioned with methanol and water. Samples were loaded, washed twice with 0.1% formic acid in water and 10% acetonitrile, and eluted with 0.1% formic acid in 80% acetonitrile. The eluate was dried using a SpeedVac and reconstituted in 50 µL of 0.1% formic acid in water.

Samples (2 µL) were injected onto a Thermo Fisher PepMap EasySpray column (75 µm x 500 mm, 1.7 μm) and separated using a 13-minute gradient with mobile phases of 0.1% formic acid in water (A) and 0.1% formic acid in 90% acetonitrile (B). The target analyte, SIL-OT, was analyzed in positive ion mode using parallel reaction monitoring (PRM). The MS² scan range was set to 200–550 m/z to minimize spectral interference, and HCD collision energy was empirically optimized. Quantification was based on peak area ratios of SIL-OT to IS, using quantifier transitions of m/z 565.7634→472.1939 (2+) for SIL-OT and 562.2548→468.6883 (2+) for the IS; additional qualifier ions were monitored for identity confirmation. Matrix-matched calibration standards were prepared in negative-control rat plasma and milk by serial dilution of SIL-OT across 12 concentration levels, with a constant CAM-OT IS spike (0.1 μg/mL; 2 μL) added to each calibrant and study sample. Matrix-matched calibration curves were generated in rat plasma and milk using a 12-point serial dilution series. The lower limit of quantification (LOQ) was 0.610 ng/mL, with a linear range extending to 10,000 ng/mL in plasma and 5,000 ng/mL in milk. Curves were fit by unweighted linear regression, with LOD and LOQ defined by signal-to-noise ratios of 3 and 10, respectively.

**Neurobehavioral assessment**

For all tasks, the rats were acclimated to the testing room at least 30 min prior to the start of testing. All assays were conducted by experimenters blinded to experimental group designations during testing, which occurred during the light phase. The order of behavioral tests was chosen to minimize the effects of stress on performance. To minimize any effect of hormonal cycling during the peripubertal period, behavior testing across experimental groups were conducted at the same time of day to control for variation in hormonal levels across the day or other circadian effects. Behavioral data were analyzed using appropriate parametric models, including Student's t-test and one-way, factorial, covariate, and repeated-measures analysis of variance (ANOVA). Simple main effects were used to dissect significant interactions. Longitudinal weight trajectory data were analyzed using two-way repeated-measures ANOVA with time and treatment as factors and litter as the repeated (subject) effect. Multiple pairwise comparisons were Bonferroni-corrected. When data did not meet univariate assumptions, non-parametric tests were used or appropriate data transformations were applied.

**RNA sequencing and analysis**

Total RNA integrity was determined using Agilent Bioanalyzer or 4200 Tapestation. Library preparation was performed with 5 to 10ug of total RNA with a Bioanalyzer RIN score greater than 8.0. Ribosomal RNA was removed by poly-A selection using Oligo-dT beads (mRNA Direct kit, Life Technologies). mRNA was then fragmented in reverse transcriptase buffer and heating to 94°C for 8 min. mRNA was reverse transcribed to yield cDNA using SuperScript III RT enzyme (Life Technologies, per manufacturer's instructions) and random hexamers. A second strand reaction was performed to yield ds-cDNA. cDNA was blunt ended, had an A base added to the 3' ends, and then had Illumina sequencing adapters ligated to the ends. Ligated fragments were then amplified for 12-15 cycles using primers incorporating unique dual index tags. Fragments were sequenced on an Illumina NovaSeq X Plus using paired end reads extending 150 bases. Basecalls and demultiplexing were performed with Illumina’s bcl2fastq software with a maximum of one mismatch in the indexing read.

Samples were prepared according to library kit manufacturer’s protocol, indexed, pooled, and sequenced on an Illumina NovaSeq 6000. Basecalls and demultiplexing were performed with Illumina’s bcl2fastq2 software. RNA-seq reads were then aligned and quantitated to the Ensembl release 101 primary assembly with an Illumina DRAGEN Bio-IT on-premise server running version 3.9.3-8 software. Gene counts were derived from the number of uniquely aligned unambiguous fragments by Subread: featureCount version 2.0.8. Isoform expression of known Ensembl transcripts were quantified with Salmon version 1.10.0. Sequencing performance was assessed for the total number of aligned reads, total number of uniquely aligned reads, and features detected. The ribosomal fraction, known junction saturation, and read distribution over known gene models were quantified with RSeQC version 5.04.

All gene counts were then imported into the R/Bioconductor package EdgeR ^1^ and TMM normalization size factors were calculated to adjust for samples for differences in library size. Ribosomal genes and genes not expressed in the smallest group size minus one samples greater than one count-per-million were excluded from further analysis. The TMM size factors and the matrix of counts were then imported into the R/Bioconductor package Limma ^2^. Weighted likelihoods based on the observed mean-variance relationship of every gene and sample were then calculated for all samples and the count matrix was transformed to moderated log 2 counts-per-million with Limma’s voomWithQualityWeights ^3^. The performance of all genes was assessed with plots of the residual standard deviation of every gene to their average log-count with a robustly fitted trend line of the residuals. Differential expression analysis was then performed to analyze for differences between conditions and the results were filtered for only those genes with Benjamini-Hochberg false-discovery rate adjusted p-values less than or equal to 0.05. For each contrast extracted with Limma, global perturbations in known Gene Ontology (GO) terms, MSigDb, and KEGG pathways were detected using the R/Bioconductor package GAGE ^4^ to test for changes in expression of the reported log 2 fold-changes reported by Limma in each term versus the background log 2 fold-changes of all genes found outside the respective term. The R/Bioconductor package heatmap3 ^5^ was used to display heatmaps across groups of samples for each GO or MSigDb term with a Benjamini-Hochberg false-discovery rate adjusted p-value less than or equal to 0.05. Perturbed KEGG pathways where the observed log 2 fold-changes of genes within the term were significantly perturbed in a single-direction versus background or in any direction compared to other genes within a given term with p-values less than or equal to 0.05 were rendered as annotated KEGG graphs with the R/Bioconductor package Pathview ^6^.

To find the most critical genes, the Limma voomWithQualityWeights transformed log 2 counts-per-million expression data was then analyzed via weighted gene correlation network analysis with the R/Bioconductor package WGCNA ^7^. Briefly, all genes were correlated across each other by Pearson correlations and clustered by expression similarity into unsigned modules using a power threshold empirically determined from the data. An eigengene was then created for each de novo cluster and its expression profile was then correlated across all coefficients of the model matrix. Because these clusters of genes were created by expression profile rather than known functional similarity, the clustered modules were given the names of random colors where grey is the only module that has any pre-existing definition of containing genes that do not cluster well with others. These de-novo clustered genes were then tested for functional enrichment of known GO terms with hypergeometric tests available in the R/Bioconductor package clusterProfiler ^8^. Significant terms with Benjamini-Hochberg adjusted p-values ≤ 0.05 were then collapsed by similarity into clusterProfiler category network plots to display the most significant terms for each module of hub genes in order to interpolate the function of each significant module. The information for all clustered genes for each module were then combined with their respective statistical significance results from Limma to determine whether or not those features were also found to be significantly differentially expressed.

**Supplementary Figure S1**

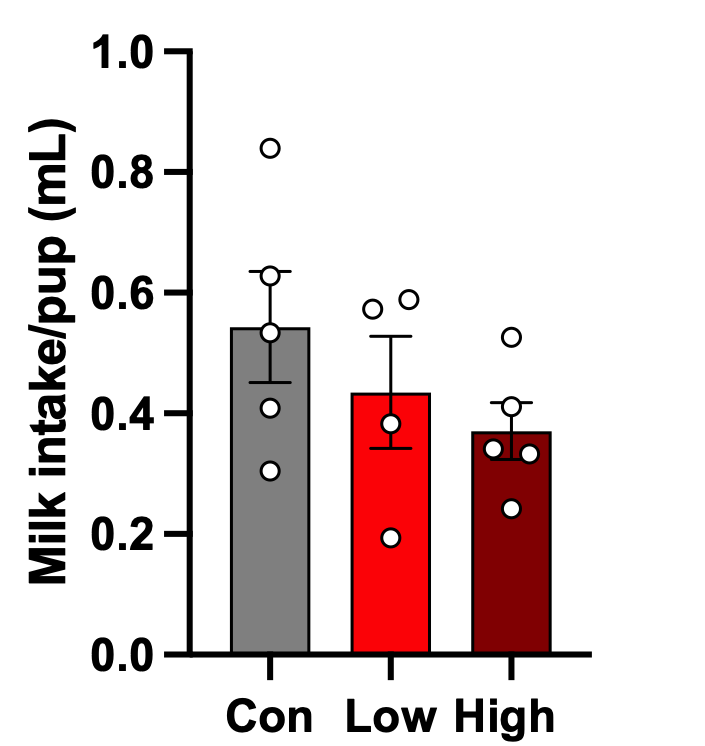

**Fig. S1.** Lactational efficiency assessed by the weigh–suckle–weigh assay in control (Con), low-dose, and high-dose OT-infused dams. Milk intake per pup showed a trend toward reduction with increasing OT dose, most pronounced in the high-dose group, though differences did not reach statistical significance. Data are presented as mean ± SEM with individual data points.

**Supplementary Figure S2**

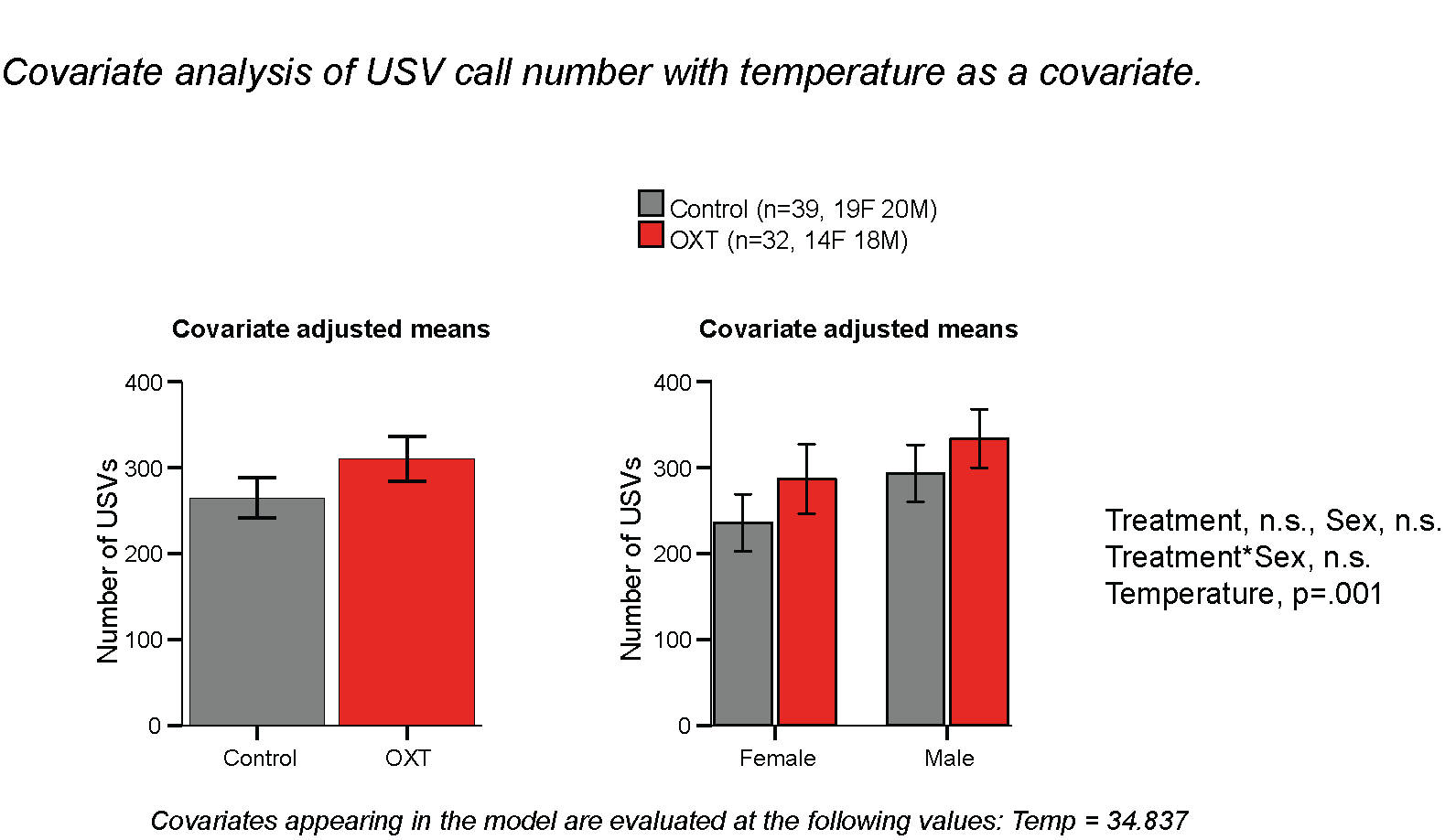

**Fig. S2.** PPH-OT increases maternal isolation-induced USVs at P8, an effect mediated by body temperature. Number of ultrasonic vocalizations (USVs) emitted during maternal isolation at postnatal day 8 in control (n = 39; 19F, 20M) and PPH-OT (OXT; n = 32; 14F, 18M) pups, shown by treatment (left) and sex (right). Covariate-adjusted means are presented with body temperature as a covariate (evaluated at 34.837°C). Treatment, sex, and their interaction were non-significant; body temperature was a significant predictor of USV number (p = 0.001), indicating that the observed increase in USVs in OXT pups was attributable to lower body temperature rather than a direct effect of oxytocin exposure. Data are mean ± SEM.

**Supplementary Figure S3**

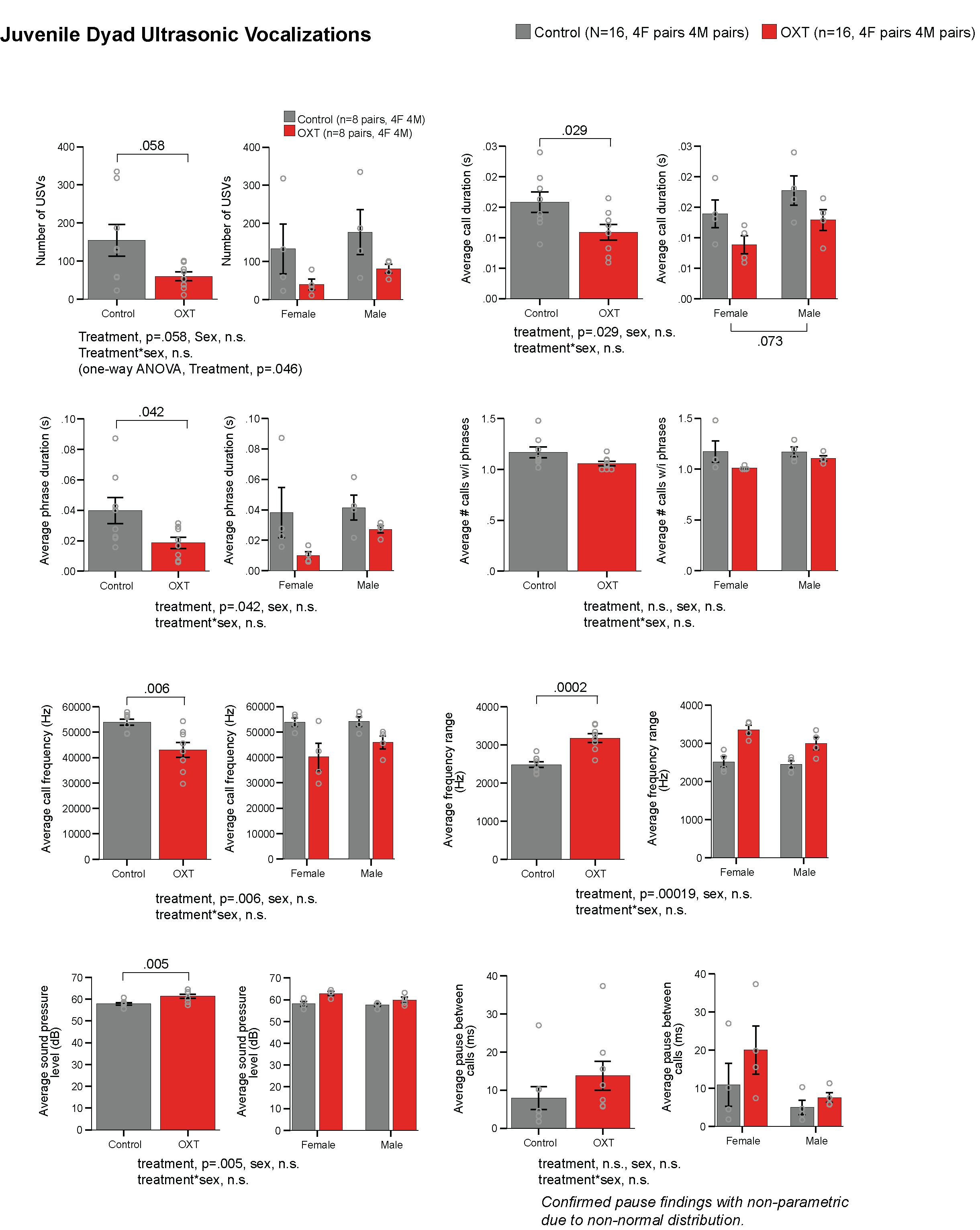

**Figure S3.** PPH-OT alters acoustic structure of juvenile dyadic USVs. Ultrasonic vocalization (USV) parameters recorded during same-group juvenile dyadic interactions in control (n = 8 pairs; 4F, 4M) and PPH-OT (OXT; n = 8 pairs; 4F, 4M) offspring, shown collapsed across sex (left panels) and by sex (right panels). OXT pairs emitted fewer total USVs (treatment p = 0.058; one-way ANOVA p = 0.046), with shorter average call duration (p = 0.029) and phrase duration (p = 0.042), lower average call frequency (p = 0.006), wider frequency range (p = 0.0002), and higher sound pressure level (p = 0.005). Average number of calls within phrases and pause between calls did not differ by treatment. No significant sex or treatment × sex interactions were detected for any measure. Data are mean ± SEM with individual data points; non-normal distributions confirmed with non-parametric tests.

**Supplementary Figure S4**

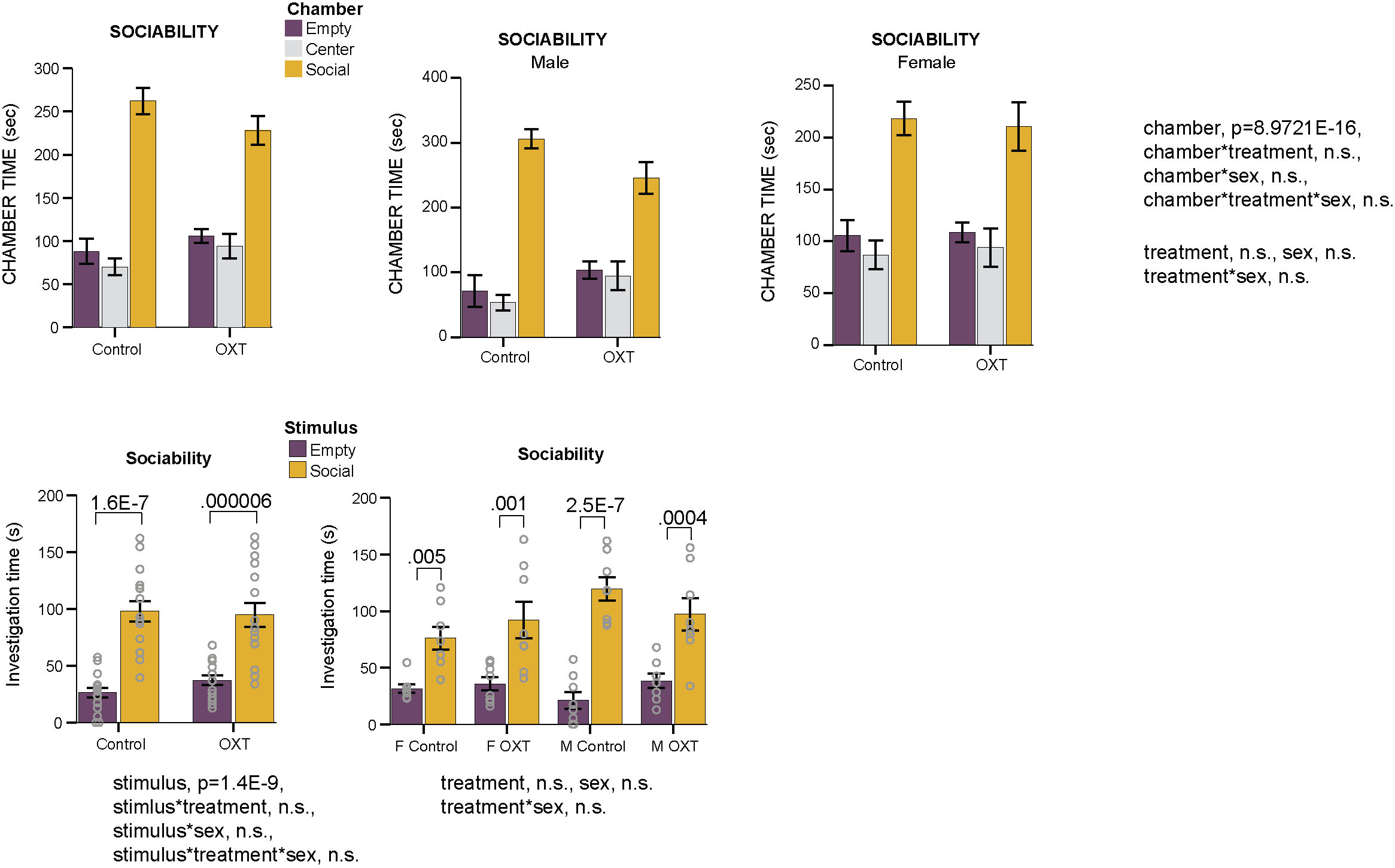

**Fig. S4.** Both control and PPH-OT offspring retain a preference for the social chamber. Time spent in each chamber (empty, center, social) during the social approach task in control and PPH-OT (OXT) offspring, shown for all animals combined (top left), males (top center), and females (top right). All groups spent significantly more time in the social chamber than the empty chamber (chamber effect, p = 8.97×10⁻¹⁶); no significant chamber × treatment, chamber × sex, or three-way interactions were detected. Investigation time directed toward the social versus empty stimulus is shown collapsed across sex (bottom left) and by sex and treatment (bottom right); all groups showed significantly greater investigation of the social stimulus (stimulus effect, p = 1.4×10⁻⁹), with no significant interactions with treatment or sex. Data are mean ± SEM with individual data points.

**Supplementary Figure S5**

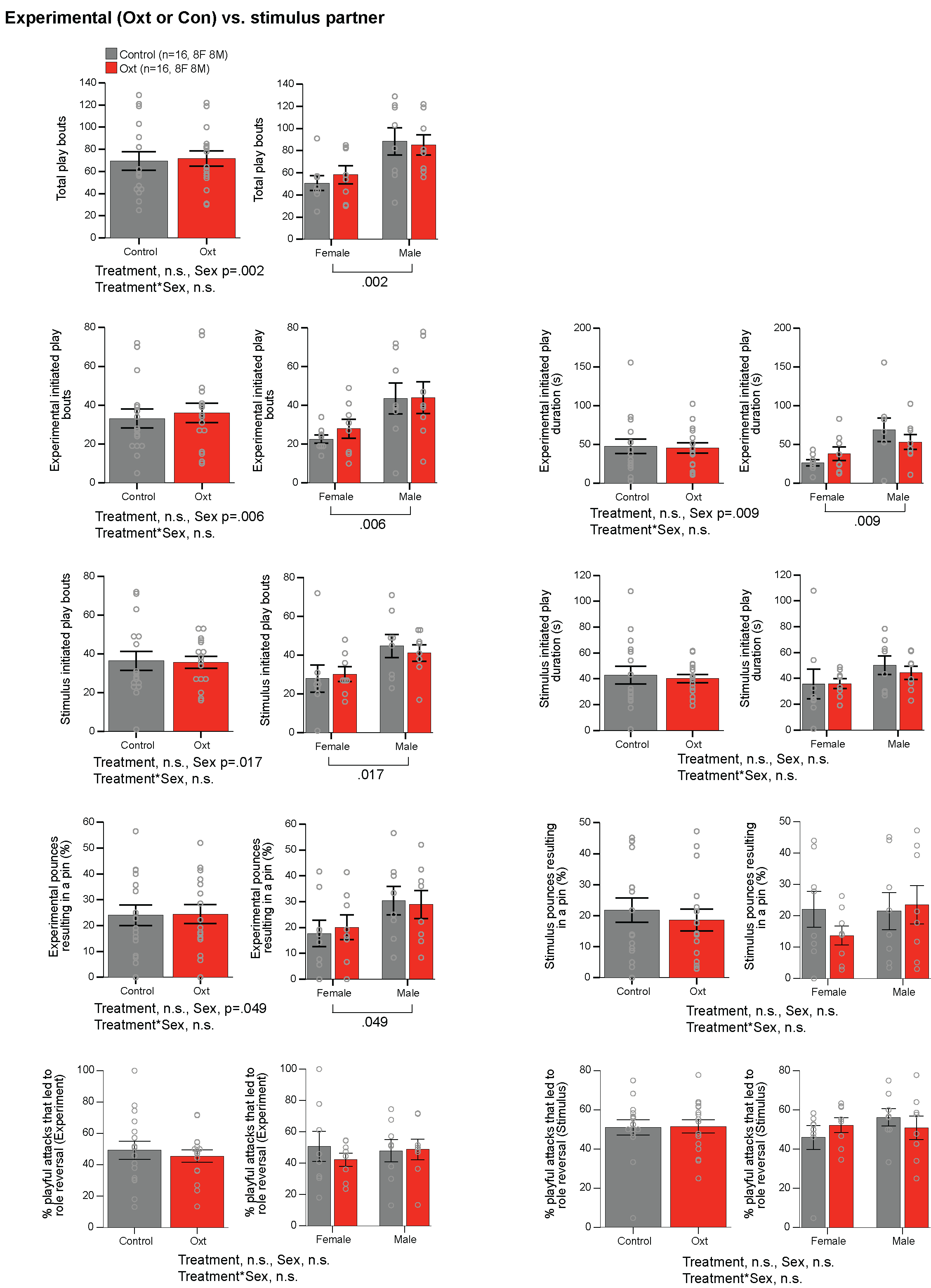

**Fig. S5.** Social play behavior is unaffected by PPH-OT but shows robust sex differences — experimental animal vs. stimulus partner pairings, bout and duration metrics. Social play metrics in control (n = 16; 8F, 8M) and PPH-OT (OXT; n = 16; 8F, 8M) experimental animals paired with same-group stimulus partners. Total play bouts, experimental- and stimulus-initiated play bouts and durations, pounces resulting in a pin, and percentage of playful attacks leading to role reversal are shown collapsed across sex (left panels) and by sex (right panels). Treatment had no significant effect on any measure. Males exhibited significantly more total play bouts (p = 0.002), experimental-initiated play bouts (p = 0.006) and duration (p = 0.009), stimulus-initiated play bouts (p = 0.017), and experimental pounces resulting in a pin (p = 0.049) than females, with no treatment × sex interactions. Data are mean ± SEM with individual data points.

**Supplementary Figure S6**

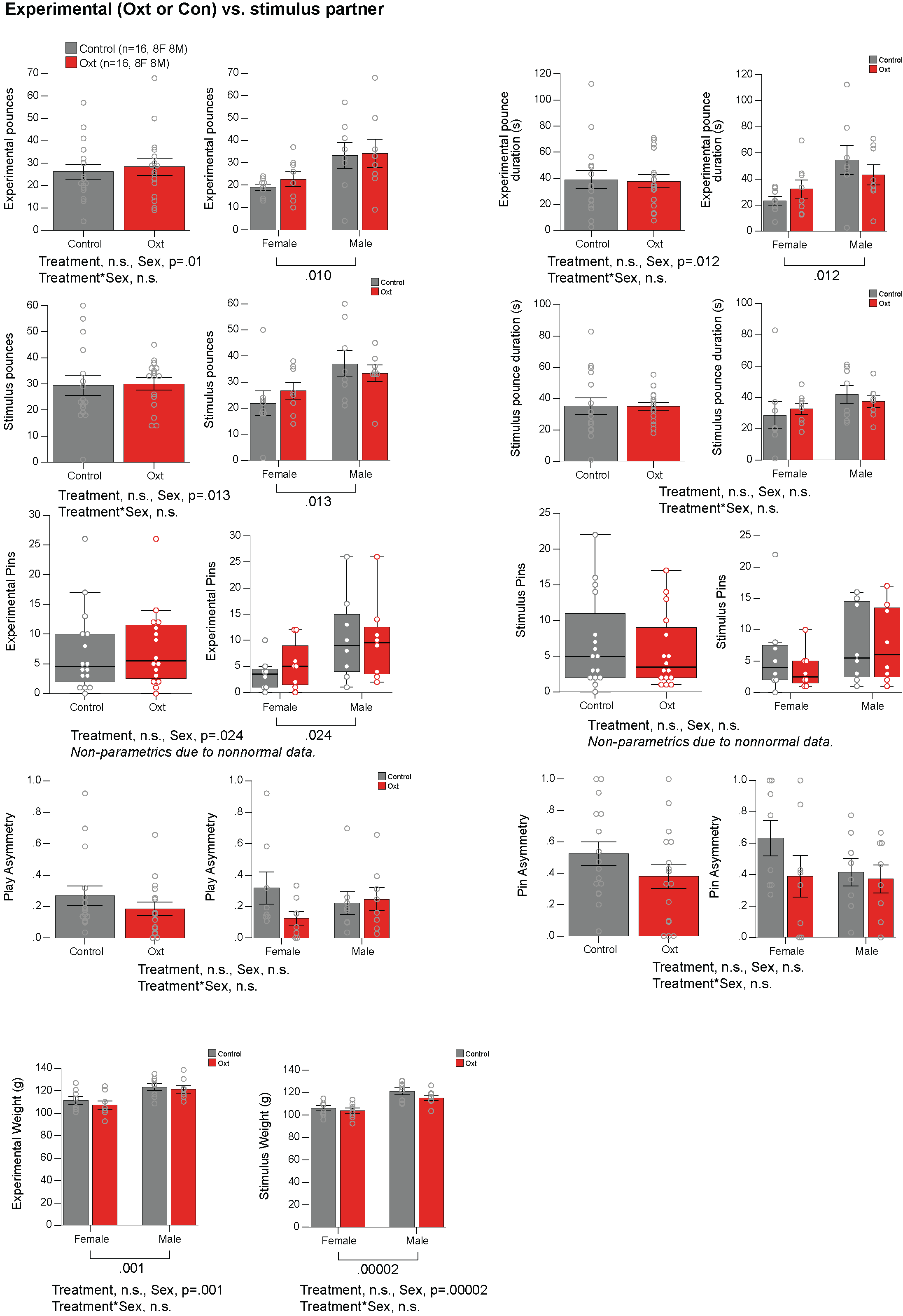

**Fig. S6.** Social play behavior is unaffected by PPH-OT but shows robust sex differences — experimental animal vs. stimulus partner pairings, pouncing, pinning, asymmetry, and body weight. Pouncing (number and duration), pinning, play asymmetry, and pin asymmetry for experimental and stimulus animals are shown collapsed across sex (left panels) and by sex (right panels) in control (n = 16; 8F, 8M) and PPH-OT (OXT; n = 16; 8F, 8M) experimental animals paired with same-group stimulus partners. Treatment had no significant effect on any measure. Males exhibited significantly more experimental pounces (p = 0.010) and longer pounce duration (p = 0.012), more stimulus pounces (p = 0.013), and more experimental pins (p = 0.024; non-parametric) than females. Play and pin asymmetry did not differ by treatment or sex. Body weight of experimental and stimulus animals differed significantly by sex (p = 0.001 and p = 0.00002, respectively) but not by treatment. Data are mean ± SEM with individual data points; non-normal distributions were analyzed with non-parametric tests.

**Supplementary Figure S7**

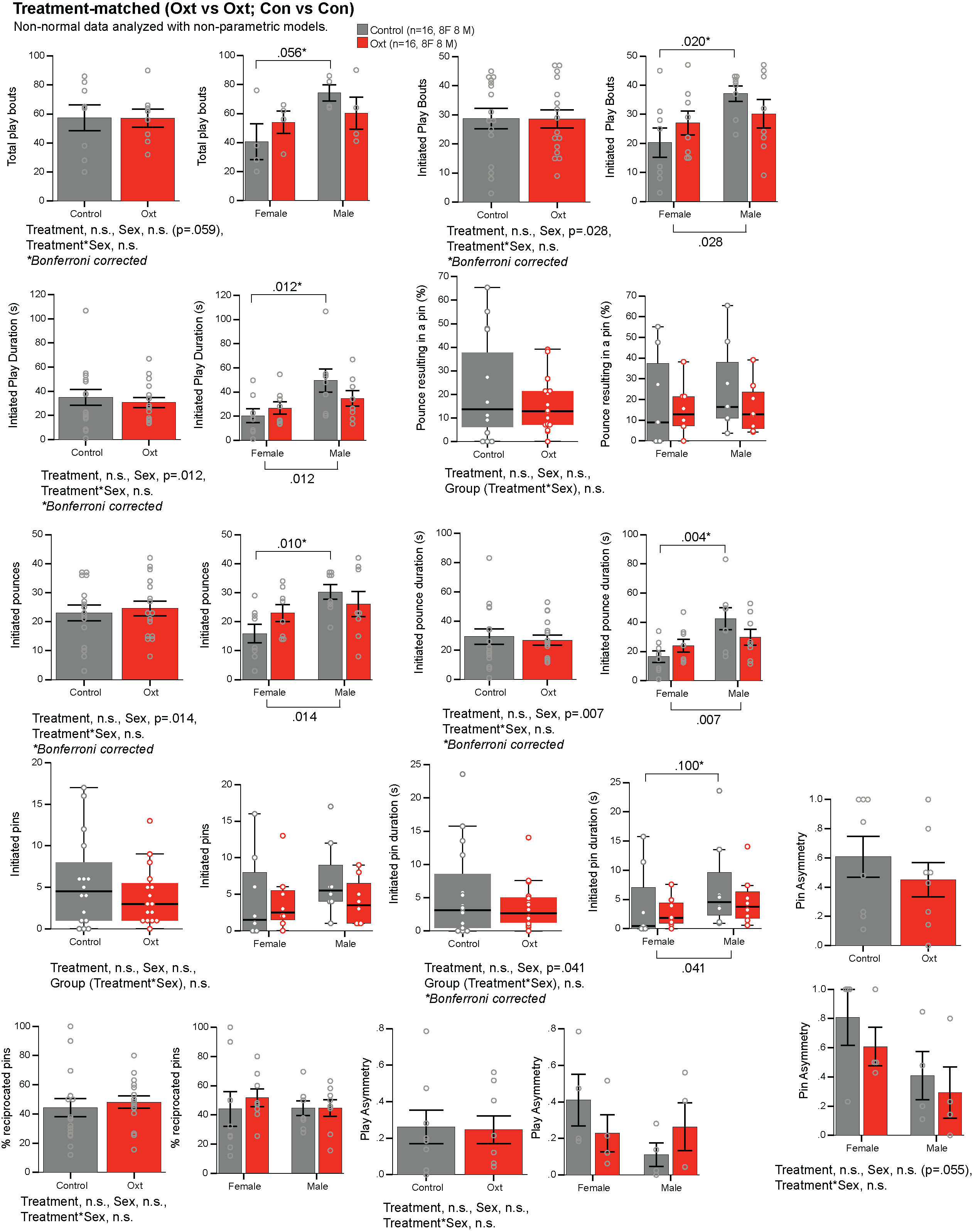

**Fig. S7.** Social play behavior is unaffected by PPH-OT but shows robust sex differences in treatment-matched pairings. Social play metrics in control (n = 16; 8F, 8M) and PPH-OT (OXT; n = 16; 8F, 8M) offspring assessed in treatment-matched dyads (Con × Con; OXT × OXT). Total play bouts, initiated play bouts and duration, initiated pounces and pounce duration, initiated pins and pin duration, percentage of reciprocated pins, and play and pin asymmetry are shown collapsed across sex (left panels) and by sex (right panels). Treatment had no significant effect on any measure. Males exhibited significantly more initiated play bouts (p = 0.028), longer initiated play duration (p = 0.012), more initiated pounces (p = 0.014) and longer pounce duration (p = 0.007), and longer initiated pin duration (p = 0.041) than females, with no treatment × sex interactions. All comparisons Bonferroni-corrected unless otherwise noted. Non-normal data were analyzed with non-parametric models. Data are median with interquartile range (box plots) or mean ± SEM (bar graphs) with individual data points.

**Supplementary Figure S8**

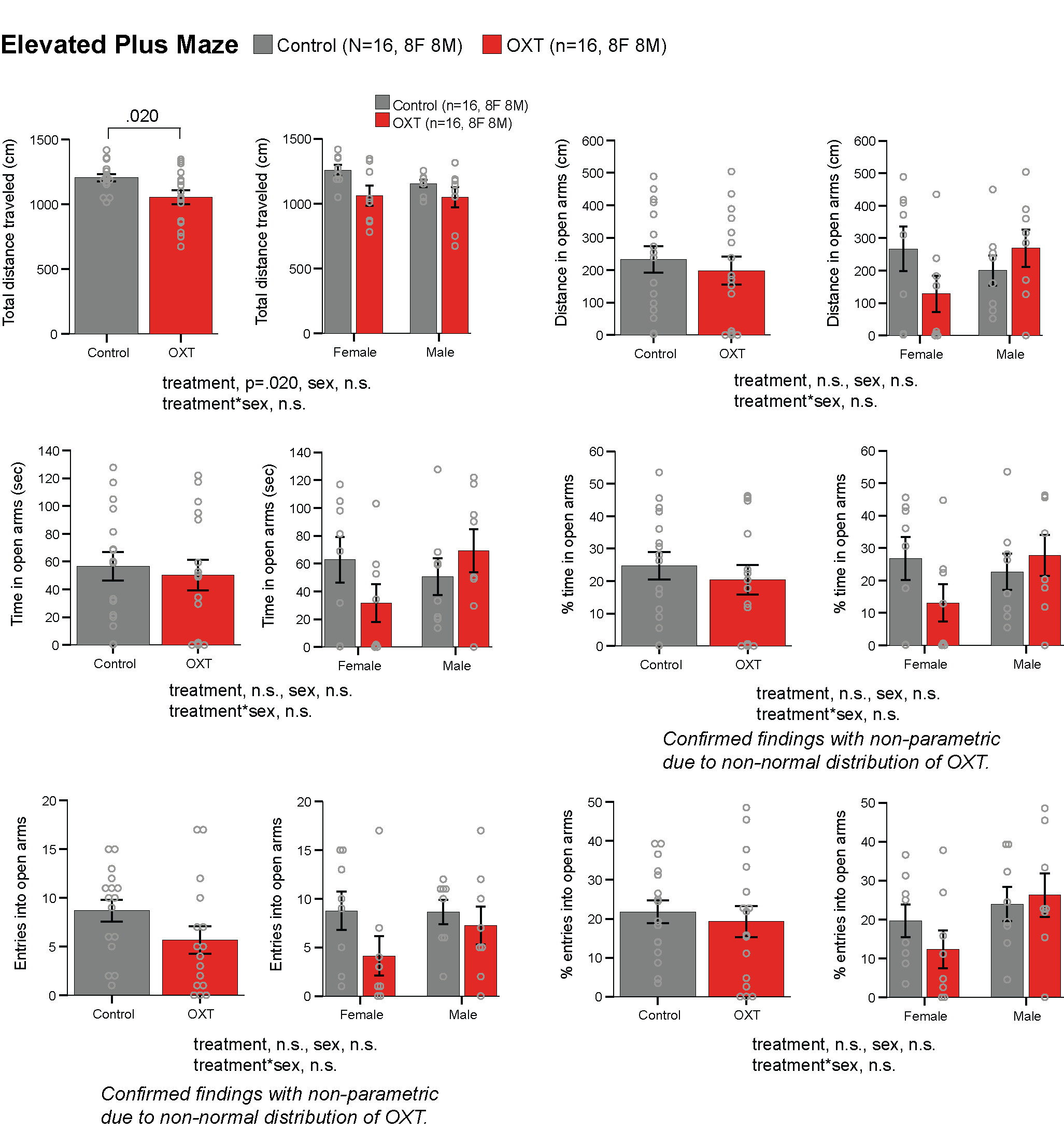

**Fig. S8.** PPH-OT reduces locomotor activity but does not affect anxiety-like behavior in the elevated plus maze. Elevated plus maze performance in control (n = 16; 8F, 8M) and PPH-OT (OXT; n = 16; 8F, 8M) offspring, shown collapsed across sex (left panels) and by sex (right panels). PPH-OT offspring traveled significantly less total distance (p = 0.020); distance traveled in the open arms, time in open arms, percentage of time in open arms, open arm entries, and percentage of open arm entries did not differ by treatment, sex, or their interaction. Non-normal distributions confirmed with non-parametric tests. Data are mean ± SEM with individual data points.

**Supplementary Figure S9**

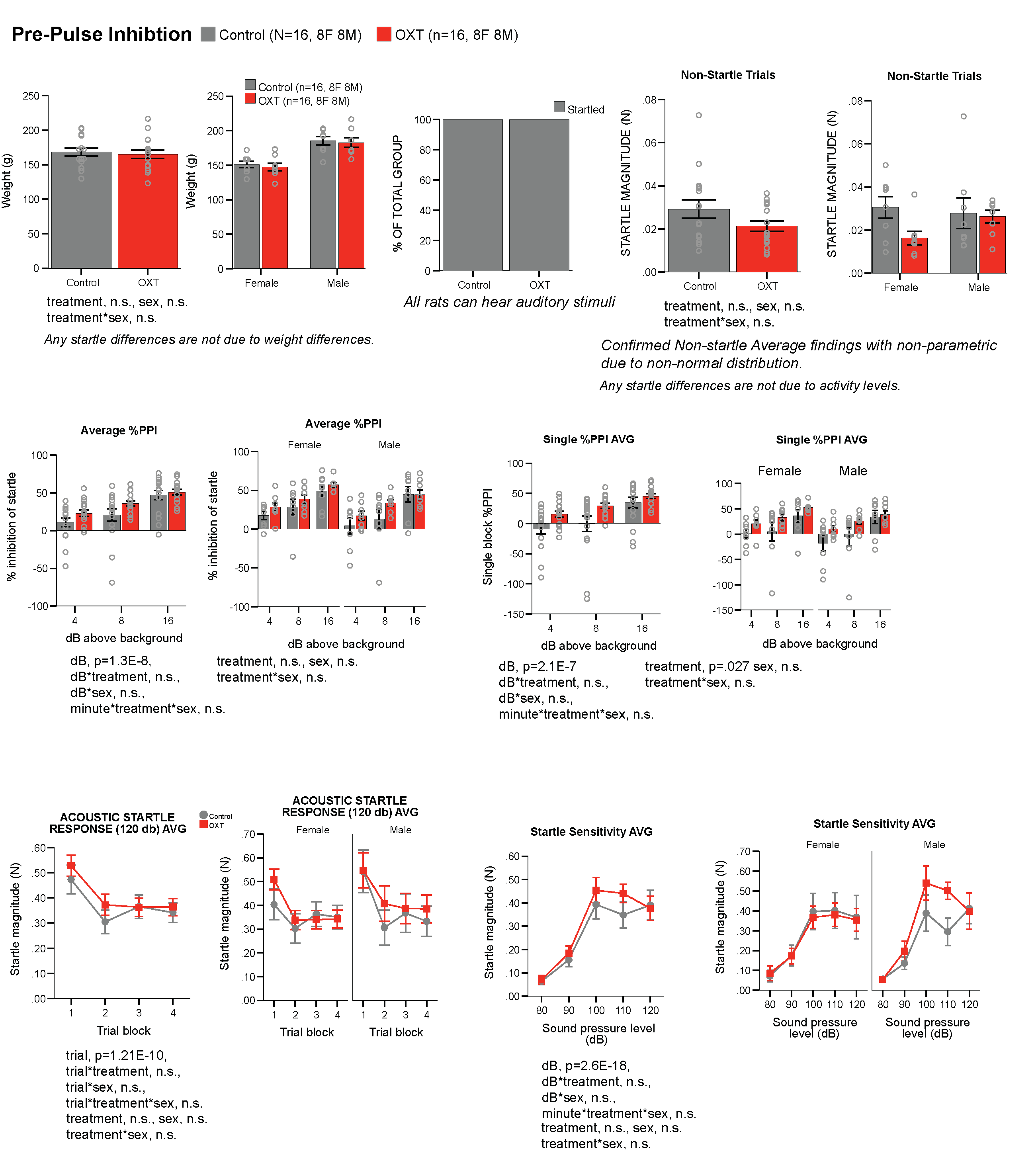

**Fig. S9.** PPH-OT does not affect sensorimotor gating assessed by prepulse inhibition. Acoustic startle and prepulse inhibition (PPI) in control (n = 16; 8F, 8M) and PPH-OT (OXT; n = 16; 8F, 8M) offspring. Body weight did not differ between groups. All animals responded to auditory stimuli; non-startle trial magnitude did not differ by treatment or sex (non-parametric). Average %PPI and single-block %PPI increased with prepulse intensity (dB effect, p = 1.3×10⁻⁸ and p = 2.1×10⁻⁷ respectively), with a significant treatment effect on single-block %PPI (p = 0.027); no treatment × dB, sex, or interaction effects were detected. Acoustic startle response habituated across trial blocks (trial effect, p = 1.2×10⁻¹⁰) and startle sensitivity increased with sound pressure level (dB effect, p = 2.6×10⁻¹⁸), with no treatment, sex, or interaction effects for either measure. Data are mean ± SEM with individual data points.

**Supplementary Figure S10**

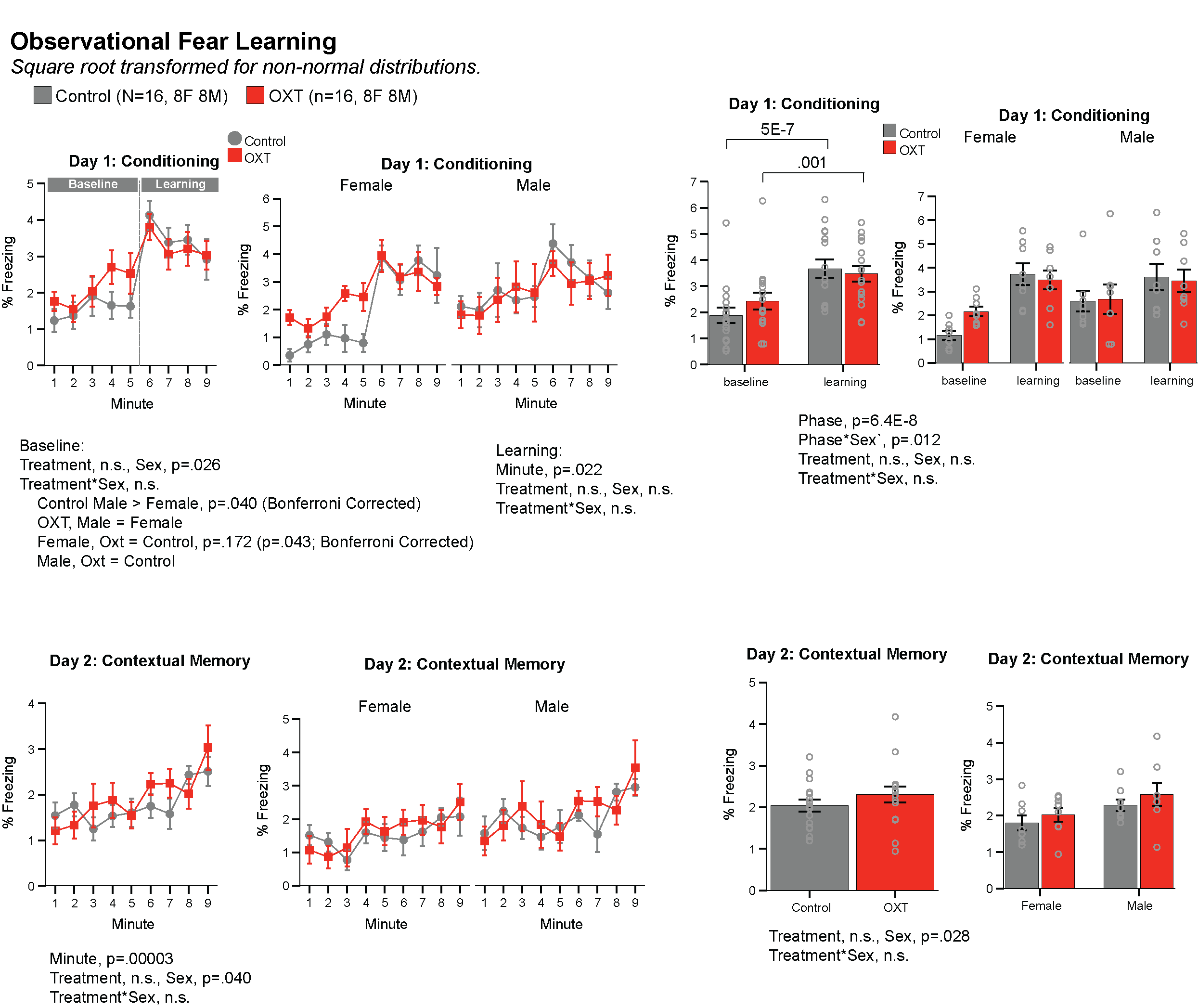

**Fig. S10.** PPH-OT does not affect observational fear learning or contextual fear memory. Percentage freezing during observational fear conditioning (Day 1) and contextual memory recall (Day 2) in control (n = 16; 8F, 8M) and PPH-OT (OXT; n = 16; 8F, 8M) offspring, shown as minute-by-minute time courses (left) and phase averages (right), collapsed across sex and by sex. On Day 1, freezing increased significantly from baseline to learning (phase effect, p = 6.4×10⁻⁸), with a phase × sex interaction (p = 0.012) driven by control males freezing more than females at baseline (p = 0.040, Bonferroni-corrected); this sex difference was absent in OXT offspring. Treatment had no significant effect on Day 1 or Day 2 freezing. On Day 2, contextual memory recall increased over time (minute effect, p = 0.00003) with a treatment × sex interaction (p = 0.040) but no main effects of treatment or sex. Data are square-root transformed; mean ± SEM with individual data points shown.

**Supplementary Table S1**

| **Group** | **Dams** | **Pups** | **Data reported** |
| --- | --- | --- | --- |
| ***Cohort 1 — dams*** | | | |
| Control dams | 5 | — | Dam-level behavioral and molecular assays (Fig. 1D–G) |
| PPH-OT dams (data contributed) | 4 | — | Dam-level behavioral and molecular assays (Fig. 1D–G) |
| PPH-OT dam excluded | 1 | — | Failed pup retrieval; reported as exclusion in Methods/Results |
| ***Cohort 1 — pups*** | | | |
| Neonatal cortical OTR qPCR (sacrificed P2) | — | 10 | Cortical OTR expression (Fig. 1I) |
| P8 maternal isolation USVs† | — | 39 | USV count and temperature (Fig. 3B) |
| Behavioral battery (P26–P49; 8M + 8F per group) | — | 32 | Weight trajectory, anxiety, social play, Y-maze, dyadic USVs, PPI, social approach, fear learning (Figs. 1H, 3C–O) |
| ***Cohort 2 — RNA-seq (Cohorts 2a + 2b combined; ≥9 independent litters per condition)*** | | | |
| Cohort 2b dedicated Con dams | 4 | — | Contributed litters for RNA-seq |
| Cohort 2b dedicated PPH-OT dams | 5 | — | Contributed litters for RNA-seq |
| RNA-seq pups — Con (10M + 10F) | — | 20 | mPFC and hypothalamus RNA-seq (Figs. 4–6) |
| RNA-seq pups — PPH-OT (10M + 10F) | — | 20 | mPFC and hypothalamus RNA-seq (Figs. 4–6) |
| ***Cohort 3 — lactational transfer of OT (separate nursing dams)*** | | | |
| OT EIA Con dams | 3 | — | Breast milk OT concentrations (Fig. 2C) |
| OT EIA PPH-OT dams | 3 | — | Breast milk OT concentrations (Fig. 2C) |
| OT EIA pups (3 pups/dam; blood pooled per litter) | — | 18 | Blood and brain samples (Fig. 2C) |
| SIL-OT LC-MS/MS Con dams | 2 | — | Milk and plasma SIL-OT quantification (Figs. 2D–H) |
| SIL-OT LC-MS/MS PPH-OT dams | 3 | — | Milk and plasma SIL-OT quantification (Figs. 2D–H) |
| SIL-OT LC-MS/MS pups (3 pups/dam; blood pooled per litter) | — | 15 | Blood and brain samples (Figs. 2D–H) |
| ***Dose-finding pilot*** | | | |
| High-dose OT dams | 5 | — | Maternal behavior and lactation at high OT dose (Supplementary File) |
| **Total dams** | **35** | **—** | **Includes 1 excluded PPH-OT dam** |
| **Total pups** | **—** | **154** | **10 (P2 qPCR) + 39 (P8 USV†) + 32 (behavioral battery) + 40 (RNA-seq) + 33 (Cohort 3)** |
| **Manuscript total** | **35** | **154** | **189 animals** |

**Table 1.** Animals used in this study. A total of 189 animals (35 dams and 154 pups) contributed to experimental outcomes reported in this manuscript. Experiments were conducted across multiple cohorts over an approximately 2-year period. Con = control (saline); PPH-OT = postpartum oxytocin (600 μg/mL, 10 μL/h × 24 h i.v. via iPrecio pump); M = male; F = female; P = postnatal day; PP = postpartum day; OTR = oxytocin receptor; USV = ultrasonic vocalization; SIL-OT = stable isotope-labeled oxytocin; mPFC = medial prefrontal cortex; PPI = prepulse inhibition. † P8 USV pups (n = 39; 23 control + 16 PPH-OT) were tested at postnatal day 8 and subsequently returned to their dams. These animals were not used in the behavioral battery reported in this study (P26–P49). The 32 pups in the behavioral battery (P26–P49) are distinct individuals selected from the same litters after litter standardization.

**Supplementary Table S2:** Timeline of behavioral assessment. The order of and age at behavioral testing with related phenotype-specific assessment in rat offspring. P: postnatal day.

| Clinical Phenotype | Specific Assessment | Behavioral Test | Age |
| --- | --- | --- | --- |
| Physical robustness | Weight measurements | -- | P8, P26 |
| Early communicative delay | Vocalization levels | Maternal isolation induced USV recordings | P8 |
| Anxiety | Approach/avoidance | Elevated plus maze | P26 |
| ASD | Social interaction behavior | Juvenile social play | P28 |
| ASD | Working memory, behavioral inflexibility | Spontaneous alternation Y-maze | P30 |
| ASD | Communication | Juvenile dyad USVs | P34 |
| ASD | Sensorimotor gating | Acoustic startle/PPI test | P41 |
| ASD | Sociability | Social approach | P43 |
| ASD | Social emotional learning (empathy-related) | Observational fear learning | P48-49 |

**Supplementary Data Files**

**Data S1**_mPFC_Oxt_v_Con_Differential_Expression_Collapsed

**Data S2**_group_mPFC_oxt_M-group_mPFC_con_M_Differential_Expression

**Data S3**_group_mPFC_oxt_F-group_mPFC_con_F_Differential_Expression

**Data S4**_Interaction_mPFC_Male_Oxt-cont_v_Female_Oxt-cont_Differential_Expression

**Data S5**_mPFC_Oxt_v_Con_GAGE_GO_Biological_Process_Results

**Data S6**_mPFC_Oxt_v_Con_GAGE_GO_Molecular_Function_Results

**Data S7**_mPFC_Oxt_v_Con_GAGE_KEGG_Signaling_and_Metabolism_Results

**Data S8**_group_mPFC_oxt_M-group_mPFC_con_M_GAGE_GO_Biological_Process_Results

**Data S9**_group_mPFC_oxt_M-group_mPFC_con_M_GAGE_GO_Molecular_Function_Results

**Data S10**_group_mPFC_oxt_M-group_mPFC_con_M_GAGE_KEGG_Signaling_and_Metabolism_Results

**Data S11**_group_mPFC_oxt_F-group_mPFC_con_F_GAGE_GO_Biological_Process_Results

**Data S12**_group_mPFC_oxt_F-group_mPFC_con_F_GAGE_GO_Molecular_Function_Results

**Data S13**_group_mPFC_oxt_F-group_mPFC_con_F_GAGE_KEGG_Signaling_and_Metabolism_Results

**Data S14**_NDD_enrichment_mPFC

**Data S15**_Hypothal_Oxt_v_Con_Differential_Expression_Collapsed

**Data S16**_group_Hypothal_oxt_M-group_Hypothal_con_M_Differential_Expression-2

**Data S17**_group_Hypothal_oxt_F-group_Hypothal_con_F_Differential_Expression

**Data S18**_Interaction_Hypothal_Male_Oxt-cont_v_Female_Oxt-cont_Differential_Expression

**Data S19**_Hypothal_Oxt_v_Con_GAGE_GO_Biological_Process_Results

**Data S20**_Hypothal_Oxt_v_Con_GAGE_GO_Molecular_Function_Results

**Data S21**_Hypothal_Oxt_v_Con_GAGE_KEGG_Signaling_and_Metabolism_Results

**Data S22**_group_Hypothal_oxt_M-group_Hypothal_con_M_GAGE_GO_Biological_Process_Results

**Data S23**_group_Hypothal_oxt_M-group_Hypothal_con_M_GAGE_GO_Molecular_Function_Results

**Data S24**_group_Hypothal_oxt_M-group_Hypothal_con_M_GAGE_KEGG_Signaling_and_Metabolism_Results

**Data S25**_group_Hypothal_oxt_F-group_Hypothal_con_F_GAGE_GO_Biological_Process_Results

**Data S26**_group_Hypothal_oxt_F-group_Hypothal_con_F_GAGE_GO_Molecular_Function_Results

**Data S27**_group_Hypothal_oxt_F-group_Hypothal_con_F_GAGE_KEGG_Signaling_and_Metabolism_Results
